# Antagonism between salicylic acid and auxin responses directs root development during *Tobamovirus* infection

**DOI:** 10.64898/2026.09.17.752527

**Authors:** Daniela Weiss, Felix Shaya, Mira Carmeli-Weissberg, Ben N. Mansfeld, Ziv Spiegelman

## Abstract

Viral symptoms in plants are structural and physiological changes resulting from virus-induced manipulation of host pathways. While the effects of viruses on above-ground organs have been well-documented, their impact on the root system remains largely understudied. In tomato (*Solanum lycopersicum*), the emerging tomato brown rugose fruit virus (ToBRFV; Genus: *Tobamovirus*) causes significant reductions in root branching. However, the mechanisms underlying this symptom remained unexplored. Here, we provide evidence that tobamovirus-mediated suppression of root branching is conserved between ToBRFV and tomato mosaic virus (ToMV), with ToMV exhibiting a more severe effect over root development. Transcriptome analysis of tobamovirus-infected roots revealed a major shift from growth to defense, marked by the early activation of salicylic acid (SA) signaling. Conversely, auxin signaling was extensively reprogrammed; For example, auxin signaling suppressors such as *Aux/IAAs* were activated while expression of auxin response factors (ARFs), was downregulated. Importantly, *SlARF19a* showed reduced expression during infection. Functional analysis of *Slarf19a* mutants confirmed its critical role in root branching and the root’s developmental response to ToMV. We conclude that tobamovirus-mediated root suppression involves an inverse relationship between SA-mediated defense and auxin-mediated growth and identify *SlARF19a* as a key regulator in this developmental trade-off.

## Introduction

Many plant viruses are detrimental pathogens, causing severe yield losses in various crops (Jones & Naidu, 2019; Panno et al., 2021; Tatineni et al., 2023). The *Tobamovirus* genus (Family: *Virgaviridae*) comprises 37 known species that infect a wide range of host plants across several families, including *Solanaceae*, *Brassicaceae*, and *Cucurbitaceae* (Adams et al., 2017; Zamfir et al., 2023). Of particular note are harmful pathogens such as Tomato mosaic virus (ToMV), Tobacco mosaic virus (TMV) and Cucumber green mottle mosaic virus, which cause dramatic reductions in yield and fruit quality of many crop species (Ishibashi et al., 2023; Jones, 2021; Spiegelman and Dinesh-Kumar, 2023). Since its recent discovery, tomato brown rugose fruit virus (ToBRFV) has caused widespread damage in commercial tomato production systems and has rapidly escalated into a global pandemic (Luria et al 2017; Hak and Spiegelman 2021; Salem et al., 2023; Zhang et al., 2022).

Tobamovirus genomes are single-stranded, positive-sense RNAs (+ssRNA) of approximately 6.4 Kb, which encode two subunits of the RNA-dependent RNA polymerase (RdRp) required for viral replication, a movement protein (MP) that enables cell-to-cell spread and a coat protein (CP) that encapsulates the viral RNA into rod-shaped particles and promote systemic movement (Heinlein, 2015; Ishibashi and Ishikawa, 2016; Ibrahim et al., 2025). As part of the infection process, some viral proteins directly interact with host proteins involved in plant growth and development to increase the efficiency of viral infection (Gnanasekaran et al., 2023; Gao et al., 2025).

Viral symptoms are defined as structural and physiological changes induced directly by the virus or its interaction with various host pathways (Gao and Lozano-Duran, 2025). These often include plant stunting, chlorosis, necrosis, fruit spotting, leaf curling, blistering, narrowing and yellowing (Pallas and Garcia, 2011). Symptom development results from a multifaceted interaction between the virus and the host plant (Pallas and Garcia, 2011; Mandadi and Scholthof, 2013; Gao and Lozano-Duran, 2025). In addition, many symptoms are attributed to the activity of silencing suppressors – viral-encoded proteins that inhibit post-transcriptional gene silencing pathways (Baulcombe, 2015). These pathways are crucial for antiviral defense while also regulating factors essential for plant growth and development in healthy plants (Kasschau et al., 2003; Kubota et al., 2003; Chellappan et al., 2005; Hendelman et al., 2013; Csorba et al., 2015). For example, the appearance of virus-associated ringspots and necrotic lesions are symptomatic manifestations of failed immune responses triggered in the infected host (Pallas and Garcia, 2011; Inaba et al., 2011).

Tobamovirus infections elicit many transcriptional and hormonal changes in the host. One such change is the induction of the phytohormone salicylic acid (SA), which triggers local immune responses as well as generates systemic acquired resistance against viruses (Malamy et al., 1990; Chivasa et al., 1997; Singh et al., 2004; Mahmoud et al., 2024; Murphy et al., 2020; Gupta et al., 2023). Apart from its role as a defense hormone, SA is a negative regulator of plant growth and development (Li et al., 2022; Pokotylo et al., 2022). Elevated levels of SA often result in plant stunting as a result of reprogramming of the plant transcriptional pathways and crosstalk with important plant growth regulators such as auxin (Pokotylo et al., 2022; Tan et al., 2020). In addition, SA dramatically affects root development by suppressing auxin signaling events that control lateral root initiation and root patterning (Pasternak et al., 2019; Bagautdinova et al., 2022). While the function of SA has been extensively explored during the plant’s antiviral immune responses, little is known as to the function of SA in virus-mediated manipulation of host development.

Tobamoviruses typically enter the plant mechanically via small wounds or scratches on the surface of leaves (Gerola et al., 1969). Upon penetration into the host cell, tobamoviruses translate, replicate and are further transported into adjacent cells through plasmodesmata intercellular channels (Heinlein, 2015; Kan and Citovsky 2025; Ibrahim et al., 2025). The virus then enters the plant vascular system to be transported long-distance through the phloem, which transports photoassimilates and information molecules from source, sugar producing organs, such as mature leaves, to sinks, sugar consuming organs such as young leaves, flowers, fruit and roots (Turgeon and Wolf, 2009; Kehr et al., 2022). The rapid accumulation of TMV in roots was already documented more than 90 years ago by Samuel (1934). Yet, while the effect of viruses on above-ground organs has been extensively explored, the impact of viruses on roots has been largely understudied.

The plant root system plays a fundamental role in development and physiology by enabling the uptake of water and nutrients and providing structural anchorage. These traits are determined by root developmental plasticity, which is continuously modified by different environmental and molecular signals (Schneider et al., 2020; Karlova et al., 2021; Dini-Andreote et al., 2025). Although more recent studies have demonstrated root invasion of viruses in different pathosystems (e.g. Valentine et al., 2002; Lunello et al., 2007; Gosalves-Bernal et al., 2008; Peltier, et al., 2011), most focus was on root crops such as sweet potato and beetroot (Peltier et al., 2011; Thiel et al., 2009; Villordon and Clark, 2014). For example, the Beet necrotic yellow vein virus p25 protein interacts with multiple Aux/IAA proteins, suppressors of the auxin response, to trigger ’rhizomania’ – a massive proliferation of lateral roots (Muelender et al., 2022). The effects of other viruses, including tobamoviruses, on root development remained to be elucidated.

Recently, we showed that ToBRFV significantly affects root growth development in tomato (Vaisman et al., 2022). ToBRFV invades the root systems early during the first week of infection in tomato plants, resulting in steep reduction in root biomass and a decrease in root-to-shoot ratio. Interestingly, ToBRFV infection resulted in marked suppression of root branching, suggesting the involvement of specific lateral root development pathways. Indeed, viral root symptoms were associated with the induction of *Auxin response factor 10a* (*SlARF10a*), a homolog of *Arabidopsis thaliana ARF10* - a known regulator of root development (Liu et al., 2007). However, mutations in *SlARF10a* were not sufficient to corroborate its effect on root branching, suggesting the involvement of other factors in virus-mediated suppression of root development (Vaisman et al., 2022).

Here, we investigated the transcriptional mechanism of tobamovirus-mediated suppression of root branching. We find that the effect on tomato root development is conserved between the tobamoviruses ToBRFV and ToMV. Transcriptome and hormone analysis of roots revealed a strong SA-mediated response in early root ToMV infection. Interestingly, application of SA alone was sufficient to emulate root responses to tobamoviruses, and ToMV-infected roots were less sensitive to SA application. These changes were associated with the reprogramming of auxin-mediated responses, including down-regulation of *SlARF19a*. Analysis of *Slarf19a* mutant reveals a key role for this gene in tomato root branching, and the root response to tobamoviruses. We conclude that the root response to tobamoviruses involves an inverse relationship between SA and auxin signaling.

## Material and methods

### Plant Materials and Growth Conditions

Tomato (*Solanum lycopersicum*) cv. Moneymaker (accession no. LA2706) seeds were obtained from the Tomato Genetics Resource Center, University of California, Davis, CA, USA. Seeds of cv. M82 (accession no. LA3475) and *arf* mutant lines (cv. M82 background), including *arf19a-1* (*Solyc07g016180*) and *arf8a* (*Solyc03g031970*) (Israeli et al., 2019; Israeli et al., 2023; Marash et al. 2024) were kindly provided by Prof. Naomi Ori (The Hebrew University of Jerusalem, Israel). All *arf* mutant lines were genotyped using gene-specific primers to confirm the presence of different mutations (listed in Supplementary table S1).

All experiments were conducted in either semi-controlled greenhouses maintained at 25 °C or growth chambers under a 16 h light / 8 h dark photoperiod at 25 °C. Seeds were sown in commercial soil (Pelemix Green 90). At the stage of fully expanded cotyledons and the emergence of the first two true leaves, seedlings were either transplanted into 8 cm^3^ pots containing supplemented with fertilizer (Ecogan Perfect, 4 ml per liter) or transferred to hydroponic systems. For transcriptome analysis, seedlings at the stage of fully expanded cotyledons were transferred directly to hydroponic systems.

For hydroponic growth, each system consisted of opaque 16 L containers equipped with lids containing 12 evenly spaced openings for plant placement. Continuous aeration was supplied using an air pump connected to an oxygen stone to ensure adequate oxygenation of the nutrient solution. For most experiments, nutrient solution was prepared by adding 5 mL of fertilizer concentrate (SHEFFA, Ct. No. Y40048; provided by Dr. Guy Tamir) to 1 L of double-distilled water, resulting in a final electric conductivity (EC) of 2.5–3.0 mS/cm. For the transcriptome assay, a Hoagland nutrient solution was used, composed of 15 mL 2 M KNO₃, 20 mL 1 M Ca(NO₃)₂, 5 mL 1 M MgSO₄, 10 mL 1 M KH₂PO₄, 1.5 mL 0.1 M EDFS, and 5 mL of a micronutrient solution containing 0.5 mM CuSO₄·5H₂O, 0.5 mM H₂MoO₄, 2 mM MnSO₄·4H₂O, 50 mM KCl, 25 mM H₃BO₃, and 2 mM ZnSO₄·7H₂O, per 5 L of double-distilled water. Separate hydroponic containers were used for each treatment group (mock, ToMV, or ToBRFV) to maintain physical separation and prevent cross-contamination between treatments.

### Tobamovirus Inoculation

Following a short acclimation period after seedling transfer, plants were mechanically inoculated with either Tomato mosaic virus (ToMV) or Tomato brown rugose fruit virus (ToBRFV). Virus inoculum was prepared from symptomatic source leaves previously stored at −20 °C. Leaves were ground in 0.01 M phosphate buffer (NaH₂PO₄) containing a small amount of carborundum powder, using a mortar and pestle. The resulting sap was applied to the cotyledons of each seedling by gentle rubbing for mechanical inoculation.

### Root Morphology Analysis

Root tissues were harvested at the indicated days post inoculation (dpi). On each sampling date, the fresh weight of the entire root system was recorded. For plants grown in soil, root systems were carefully removed from the pots and thoroughly washed with tap water to eliminate residual substrate. Clean roots were placed in shallow trays containing a minimal volume of water and scanned using a flatbed scanner adapted for root imaging. Root morphology was quantified using WinRHIZO Pro software (Regent Instruments, Quebec, Canada) as described by Pang et al. (2011). Grayscale contrast settings were optimized for each image to enhance root detection accuracy. Quantified parameters included total root length (cm), number of forks (lateral branches), number of root tips, average root diameter (mm), surface area (cm²), and root system volume (cm³). Following image acquisition, root samples were oven-dried at 60 °C for six days to determine dry weight.

### Exogenous Salicylic Acid Application

Salicylic acid (SA; Sigma-Aldrich, Cat. No. 69-72-7) was dissolved in double-distilled water (DDW) to final concentrations of 0.5 mM, 1 mM, and 2 mM. The SA solutions were applied to tomato plants by foliar spraying until visible leaf run-off was observed. The first application was performed 5 days prior to ToMV inoculation, and the second on the day of inoculation. Additional applications were carried out at 4–5-day intervals until tissue collection, which occurred 19 days post inoculation (dpi).

### RNA Extraction, Library Preparation, and Sequencing

Total RNA was extracted from 100 mg of root tissue using the Total RNA Purification Kit (Norgen, Cat. No. 25800) according to the manufacturer’s instructions. RNA samples were shipped to Novogene (Beijing, China) for quality control and illumina TruSeq library preparation Sequencing was performed on an Illumina NovaSeq 6000 platform, generating 2 × 150 bp paired-end reads at a depth of approximately 20 million read pairs per sample. Raw reads were processed to remove adapter sequences, reads containing poly-N, and low-quality reads, resulting in high-quality clean reads for downstream analyses.

### Transcriptome analysis

For generating transcript level read counts, first the cDNA fasta file from the tomato ITAG4.0 annotation (Hosmani et al., 2019) was concatenated to the whole tomato genome file to prepare an alignment decoy file and index (https://combine-lab.github.io/alevin-tutorial/2019/selective-alignment/). RNAseq reads were then pseudo-aligned to the ITAG4.0 transcriptome using Salmon v1.5.2 (Patro et al., 2017) default settings. Read count data were imported into R using the tximport package (Soneson et al., 2016). Multiple libraries for a sample (i.e. sequencing runs) were merged using the ‘collapseReplicates’ function from the DESeq2 package (Love et al., 2014). Differential expression on the gene-level integer count values was performed using DESeq2 (Love et al., 2014). Genes with a mean of less than 20 reads across all samples were excluded from analysis. One sample from the ToBRFV inoculated plants was removed due to extremely low mapping rates (99% unmapped reads) likely caused by contamination with another sample at the sequencer. Principal component analyses were performed using the variance stabilized transformed read counts DESeq2 and a modified version of the plotPCA function from DESeq2. Differential expression was performed using “apeglm” (Zhu et al., 2019) as the Log Fold Change Shrinkage method. Genes were defined as being significantly differentially expressed if they had an adjusted p-value of less than 0.05. Normalized counts were plotted using ggplot and tidyverse (Wickham et al., 2019) functions in R. The Gene Ontology term (GO term) database for tomato, was previously created by GOMAP (Wimalanathan and Lawrence-Dill, 2021) and downloaded from doi.org/10.25739/zh2v-4p15. Gene Ontology (GO) term enrichment analysis was performed using the TopGO R package (Alexa et al., 2006) with the entire set of expressed genes set as background. Terms were considered enriched if they passed a p-value of 0.05 on the Fisher test with the “weight01” algorithm and a minimum node size of 100 genes. GO-terms were then clustered by Euclidean distances by similarity measures and reduced to parent terms using the rrvgo R package (Sayols, 2023).

### Reverse transcription quantitative PCR (RT-qPCR)

Gene expression and Tobamovirus infection were quantified by qPCR using tissue collected from young leaves (>1 cm from the shoot apex) or roots (gene expression analysis was performed on roots only). Total RNA was extracted from 50–100 mg of tissue using the Plant Total RNA Mini Kit (Geneaid, RPD300), which includes on-column DNase treatment to remove genomic DNA. For gene expression assays, an additional DNase I treatment was performed using DNase I (Thermo Fisher Scientific) according to the manufacturer’s instructions. A total of 400–500 ng of RNA was used for cDNA synthesis with the AzuraQuant II cDNA Synthesis Kit (Azura Genetics, AZ-2504), following the manufacturer’s protocol. Quantitative real-time PCR was conducted using AzuraView GreenFast qPCR Blue Mix LR (Azura Genetics, AZ-2305) in a 10 µL reaction volume (half reaction). Primer concentrations were either 150 nM or 250 nM. Reactions were run on an Applied Biosystems QuantStudio 3 Real-Time PCR Instrument (Thermo Fisher Scientific) with the following cycling conditions: 95 °C for 2 min, followed by 40 cycles of 95 °C for 5 s and 55 °C or 60 °C for 30 s, with a final step at 60 °C for melt curve analysis. Fluorescence data were analyzed using QuantStudio Design and Analysis Software v1.5.2. Relative ToMV and ToBRFV RNA levels were normalized to the reference transcript *TIP41* (*Solyc10g049850*) (Lacerda et al., 2015). Endogenous gene expression levels were calculated using the 2^–ΔΔCt method (Livak and Schmittgen, 2001). Expression values were normalized to the geometric mean of two reference transcripts, *TIP41* (*Solyc10g049850*) and *ACTIN* (*Solyc03g078400*), and were expressed relative to mock-treated control samples. A list of all primers used in this study is provided in Supplementary table S1.

### Plant Hormone Extraction and LC–MS Analysis

Plant hormone extraction was performed using a solvent-based protocol optimized for the stabilization of phytohormones. Immediately after excision, tissue samples were flash-frozen in liquid nitrogen to prevent hormone degradation. Approximately 200 mg of frozen tissue was weighed prior to extraction and ground to a fine powder in liquid nitrogen using a pre-chilled mortar and pestle. The powdered tissue was transferred to 2 mL microcentrifuge tubes containing 1 mL extraction solvent consisting of isopropanol:methanol:glacial acetic acid (79:20:1, v/v/v) and a mixture of isotope-labeled internal standards (20 ng of each). Samples were incubated with continuous vortexing for 60 min at 4 °C and centrifuged at 14,000 rpm for 15 min at 4 °C. The supernatant was transferred to a fresh tube, and the pellet was re-extracted twice by adding 0.5 mL extraction solvent, followed by vortexing, centrifugation, and collection of the supernatant. All supernatants were combined and evaporated to dryness (or to a minimal volume) at room temperature using a SpeedVac concentrator. The dried extracts were dissolved in 200 µL of 50% methanol, centrifuged at 14,000 rpm for 15 min at 4 °C, and filtered through 0.22 µm PVDF syringe filters (13 mm). Samples were stored at −20 °C until analysis. For LC–MS analysis, 5 µL of each sample was injected into the instrument. Quantification was performed for salicylic acid (SA), free auxins [indole-3-acetic acid (IAA) and indole-3-butyric acid (IBA)], conjugated auxins (IAA-Asp, IAA-Glu, IBA-Glu) and oxidized auxin (ox-IAA). Hormone levels were normalized based on the signal of the corresponding internal standard and the initial fresh weight of the tissue to allow for accurate absolute quantification across samples.

### Confocal Imaging and Fluorescence Quantification

The expression pattern of the *pDR5:VENUS* auxin reporter line (*cv.* M82, provided by Prof. Idan Efroni) was visualized using a confocal laser scanning microscope (Olympus IX 81; Fluoview 500 software) equipped with an OBIS laser line and a 20× objective. VENUS was excited at 488 nm and imaged using an BA505-525 nm emission filter. Tissue preparation and clearing was performed using the ClearSee method as described by Kurihara et al. (2015). Roots were harvested and immediately fixed in 4% paraformaldehyde (PFA; P6148, Sigma-Aldrich) in 1× phosphate-buffered saline (PBS; pH 7) for 1 h at room temperature with gentle shaking. Fixed tissues were washed twice for 1 min each in 1× PBS to remove residual fixative. Following fixation, samples were transferred to ClearSee solution (10% [w/v] xylitol, 15% [w/v] sodium deoxycholate, and 25% [w/v] urea) and cleared overnight at room temperature. Fluorescence signal intensity was quantified using ImageJ software (Shihan et al., 2021). Fluorescence measurements were done using ImageJ. For each sample, fluorescence was normalized by subtracting the mean background fluorescence from the VENUS fluorescence signal to minimize background noise and allow for accurate comparisons between samples.

## Results

### Comparative analysis of tobamovirus infection impact on tomato root development

To understand if the impact of different tobamoviruses on tomato root development is conserved, we examined the effects of two tobamovirus species, ToMV and ToBRFV on root morphology. Tomato seedlings (cv. Moneymaker) were mechanically inoculated with either ToMV or ToBRFV. Successful infection was verified in all inoculated plants using RT-qPCR on systemic leaves (Supplementary Fig. S1). Mock-inoculated plants served as controls. Roots of soil-grown plants were monitored at 7-, 14-, and 21-days post inoculation (dpi) for structural analysis (Fig. 1A). Root development was evaluated by measuring dry biomass (Fig. 1B) and morphological analyses were performed using the WinRhizo image analysis system (Fig. 1C-G). At 7 dpi, only minimal changes in root diameter of ToBRFV-infected plants and (Fig. 1E) and branch number in ToMV-infected plants (Fig. 1F) were observed. However, at 14 and 21 dpi, significant reductions were observed in both ToMV- and ToBRFV-infected root dry weight (Fig. 1B), total root length (Fig. 1C), roots volume (Fig. 1D), average diameter (Fig. 1E), number of branches (Fig. 1F), and branch density (Fig. 1G). While both ToMV and ToBRFV resulted in similar effects on root, ToMV had a significantly stronger effect over root development (Fig. 1B-G). These results suggest that different tobamoviruses have similar effects on root development by reducing root growth and branching.

**Figure 1.**
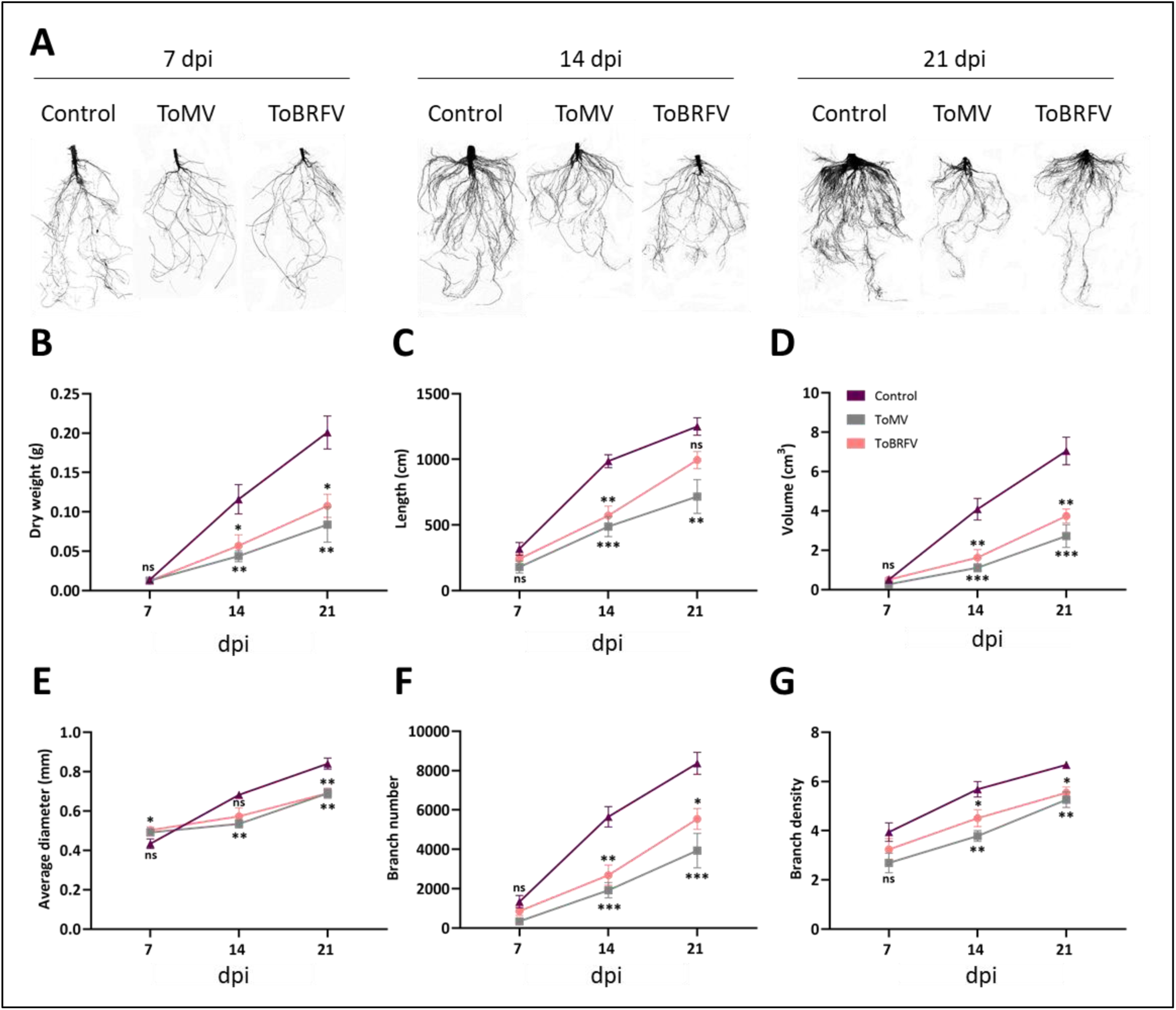
Effect of Tobamoviruses, ToMV and ToBRFV, on tomato root system development. (A) Representative scanned root systems of tomato plants inoculated with mock, ToMV, ToBRFV at 7-, 14-, and 21-days post inoculation (dpi). (B-G) Quantification of morphological root traits, including root dry weight (B) total root length (C) root volume (D) average root diameter (E) branches number (F) and root branch density (branches per cm root length) (G). Statistical analysis was performed using one-way ANOVA followed by Tukey’s post hoc test for multiple comparisons, or Dunnett’s test when variances were unequal. Data are presented as mean ± SEM. *\* P<0.05, ** P<0.01*, ***P<0.001, n= 6-7.

Molecular analysis of root development during infection requires a soil-free, non-invasive system. To do so, we constructed a hydroponic growth system, in which the roots of tomato plants (cv. Moneymaker) are easily observed as the infection progresses (Supplementary Fig. S2A). To verify the hydroponic system did not impact root symptoms we performed a second experiment in which 10-day-old seedlings were mechanically inoculated with ToBRFV or ToMV, and roots of infected plants were harvested and evaluated using Winrhizo at 10 dpi (Supplementary Fig. S2B). Consistent with soil-grown roots findings, clear suppression of root branching was already observed at 10 days post inoculation, with ToBRFV causing a milder effect as compared to ToMV (Supplementary Fig. S2, B-F): A reduction of 40.3% and 69.9% was observed in total root lengths of ToBRFV- and ToMV-infected plants, respectively (Supplementary Fig. S2C). Primary root lengths were only reduced by 19.6% and 38.7% (Supplementary Fig. S2D). A reduction of 51.5% and 78.3% in root branching was observed in ToBRFV- and ToMV-infected plants (Supplementary Fig. S2E).

Root branching density (branches per cm) decreased by 42.7% and 66.7% in ToBRFV- and ToMV-infected plants, respectively (Supplementary Fig. S2F). Collectively, these analyses established that root growth response to different tobamoviruses is conserved and can be largely attributed to suppression of root branching, with ToBRFV inducing a milder effect over root development than ToMV.

To determine the association between virus accumulation in roots and their development, RNA was extracted from hydroponically grown roots at various dpi (Fig. 2). Consistent with our previous findings (Vaisman et al., 2022), accumulation of viral RNA was evident for both viruses at 4 dpi (Fig. 2A-B). The highest accumulation of ToMV was documented at 10 dpi, remained high at 14 dpi and declined at 21 dpi (Fig. 2A), while ToBRFV RNA levels peaked at 14 dpi (Fig. 2B). These results are consistent with our previous observations that ToBRFV systemic spread is delayed compared to ToMV (Hak and Spiegelman, 2021).

**Figure 2.**
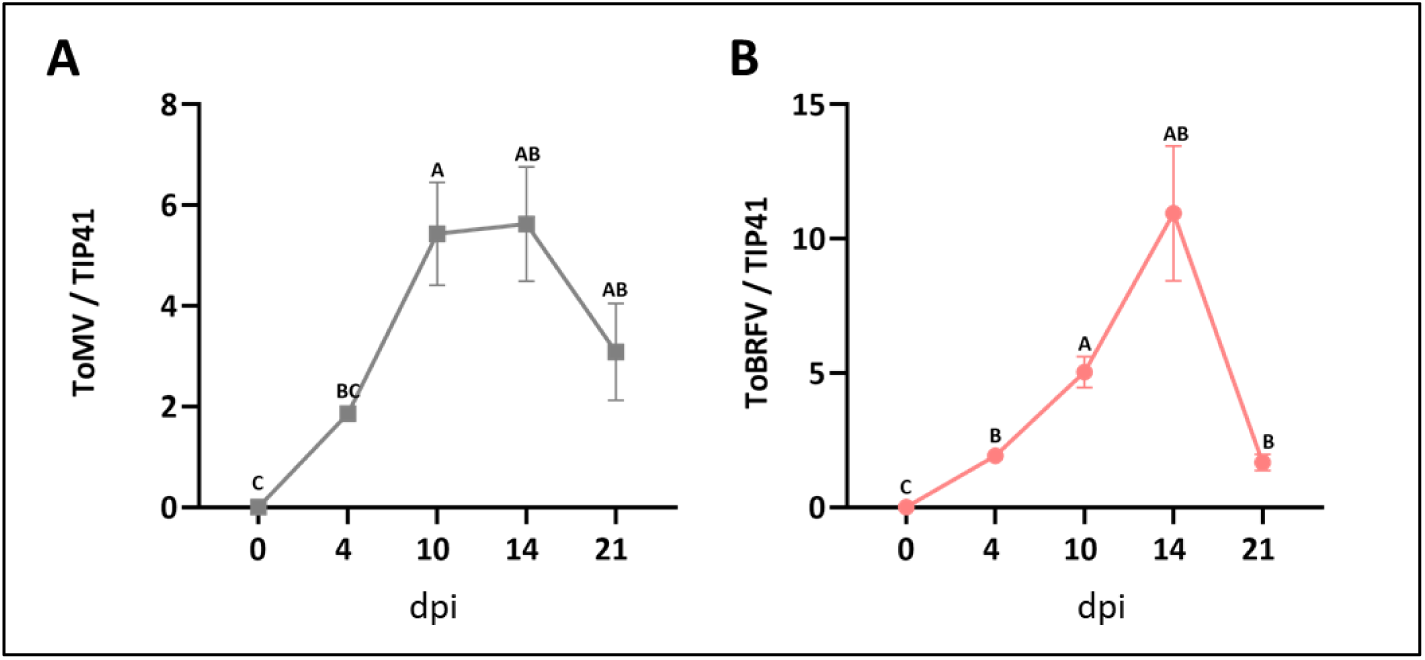
Tobamovirus viral load in roots over the course of infection. (A) ToMV and (B) ToBRFV accumulation was quantified by qPCR at 4, 10, 14, and 21 dpi. Expression levels were normalized to the housekeeping gene *TIP41*. Data represent mean ± SE of biological replicates (*n* = 8–12). Different letters indicate significant differences in Dunnett’s multiple-comparison test, or by the Kruskal–Wallis non-parametric test followed by Dunn’s multiple-comparison test (*p* < 0.05).

### Transcriptome analysis of tobamovirus-infected roots

To gain insights into the molecular mechanisms underlying tomato root responses to tobamovirus infection, transcriptome analysis was performed on roots of tobamovirus-infected tomato plants. Roots were harvested at 10 dpi, an early stage of infection at which viruses accumulate in the root system (Fig. 2) and changes in root development are already evident (Fig. 1; Supplementary Fig. S2). Principal component analysis (PCA) was performed to assess the variation within and between each treatment group. This analysis revealed that above ground inoculation with ToMV affected the overall root transcription profile shifting samples along principal component 1, which explained 56% of the variance in the data (Fig. 3A). Conversely, the effect of ToBRFV on the root transcriptome was highly variable as compared to ToMV (Fig. 3A). These results suggest that at 10 DPI, the effects of ToMV on root transcription are stronger and more consistent when compared to ToBRFV.

**Figure 3.**
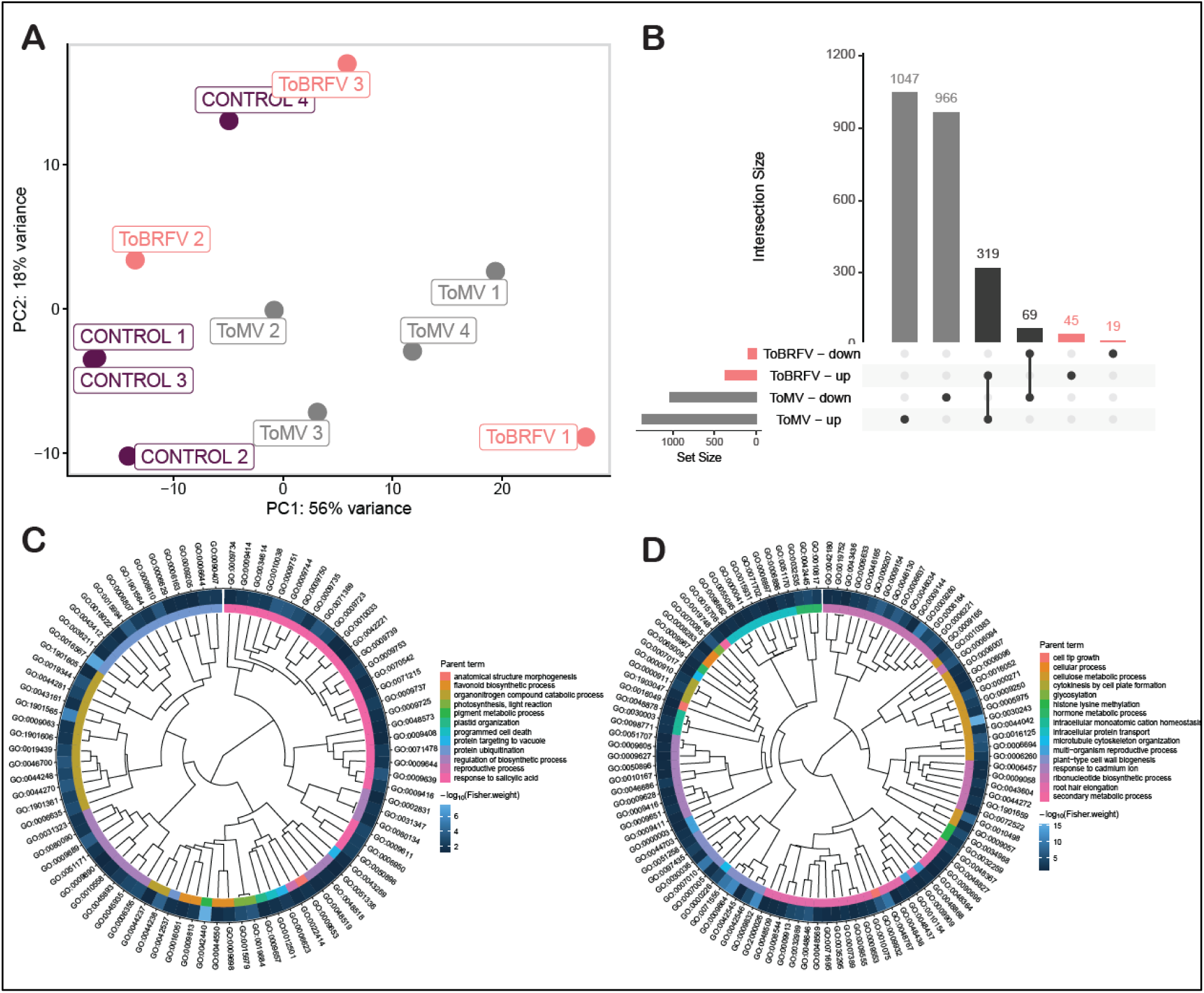
Analysis of transcriptome data from roots systems ToMV- and ToBRFV-infected tomato plants at 10 dpi. (A) Principal component analysis of the transcriptome data. (B) Differential expression analysis of up- and down-regulated transcripts in ToMV-, and ToBRFV-infected root systems. (C) Enriched GO categories in up-regulated transcripts of ToMV infected roots. (D) Enriched GO categories in down-regulated transcripts of ToMV infected roots. Terms are clustered by the Euclidian distance of their semantic similarity. Each cluster is assigned a parent term (internal color band). Color of tile for each individual GO-term indicates the -log_10_(Fisher.weight01) *p-value*.

Differential expression (DE) analysis (Fig. 3B; Supplementary table S2) revealed that 1,366 transcripts were upregulated in ToMV-infected roots but only 364 genes were upregulated in ToBRFV-infected roots (Fig. 3B). Among those genes, 319 were upregulated in both ToMV- and ToBRFV-infected root systems (Fig. 3B). In ToMV-infected roots, 1,035 transcripts were downregulated, and 88 were downregulated in ToBRFV-infected roots as compared to roots of non-inoculated plants (Fig. 3B). Of those, 69 were downregulated in both ToMV- and ToBRFV-infected roots (Fig. 3B). Gene ontology (GO) enrichment analysis revealed that transcripts upregulated by both tobamoviruses include categories related to response to stress and stimulus as well as primary and specialized metabolic processes (Supplementary table S3). Since ToMV elicited a stronger transcriptional response with more significant DEGs, we performed GO clustering analysis specifically on the ToMV set. After clustering GO terms by semantic similarity, enriched terms associated with response to salicylic acid formed the largest cluster in ToMV-infected roots (Fig. 3C).

In marked contrast, and consistent with the observed root phenotypes, downregulated transcripts were enriched for various terms related to growth and development including cell tip growth, root hair elongation, cell wall structure, and cytoskeleton organization and dynamics (Fig. 3D; Supplementary table S4). These results suggest that tobamovirus infection results in a transcriptional shift between growth and defense, affecting root architecture.

### Salicylic acid signaling is activated in tobamovirus-infected roots

As ‘response to salicylic acid’ was of the most important GO-term parent clusters we assessed the gene expression of genes in this term. Indeed, at 10 dpi, many SA-regulated transcripts were up-regulated in ToBRFV, and to a greater extent in ToMV-infected root systems (Fig. 4A). For example, the canonical defense response *Pathogenesis Related* (PR) *5* family gene *PR5-x* (*Solyc08g080620*) showed a roughly 200x increase in expression in ToMV infected roots (Fig. 4B). Since the transcriptional and root phenotype responses were stronger and more consistent in ToMV infections, we furthered our investigations using this virus. First, we quantified SA levels in ToMV-infected roots using LC-MS at 4, 10, 14 and 21 DPI (Fig. 4C). Interestingly, a dramatic spike in SA levels was observed at 4 DPI, and a decrease was observed later during infection (Fig. 4C). To understand the dynamic changes in SA signaling, the transcript abundance of SA regulatory and signaling factors was quantified at different time points throughout infection using RT-qPCR (Fig. 4D-F). Upregulation of the SA regulatory enzyme salicylic acid 5-hydroxylase (*SlS5H*) occurred at 4 dpi, consistent with the increase in SA accumulation (Fig. 4E). In addition, we assessed the expression of the SA-responsive transcripts *SlPR1a* and *SlPR-5x* during ToMV infection and observed that these are induced early and show reduction by 14 dpi, corroborating the 10 dpi RNAseq data (Fig. 4E-F). Moreover, levels of the SA receptor *non-expressor of PR genes 1* (*SlNPR1*) were also induced at 4 and 10 DPI (Fig. 4G). These results suggest that tobamovirus infection leads to an early, strong, but transient, increase in SA, resulting in SA-mediated signaling and downstream responses during the first days of infection.

**Figure 4.**
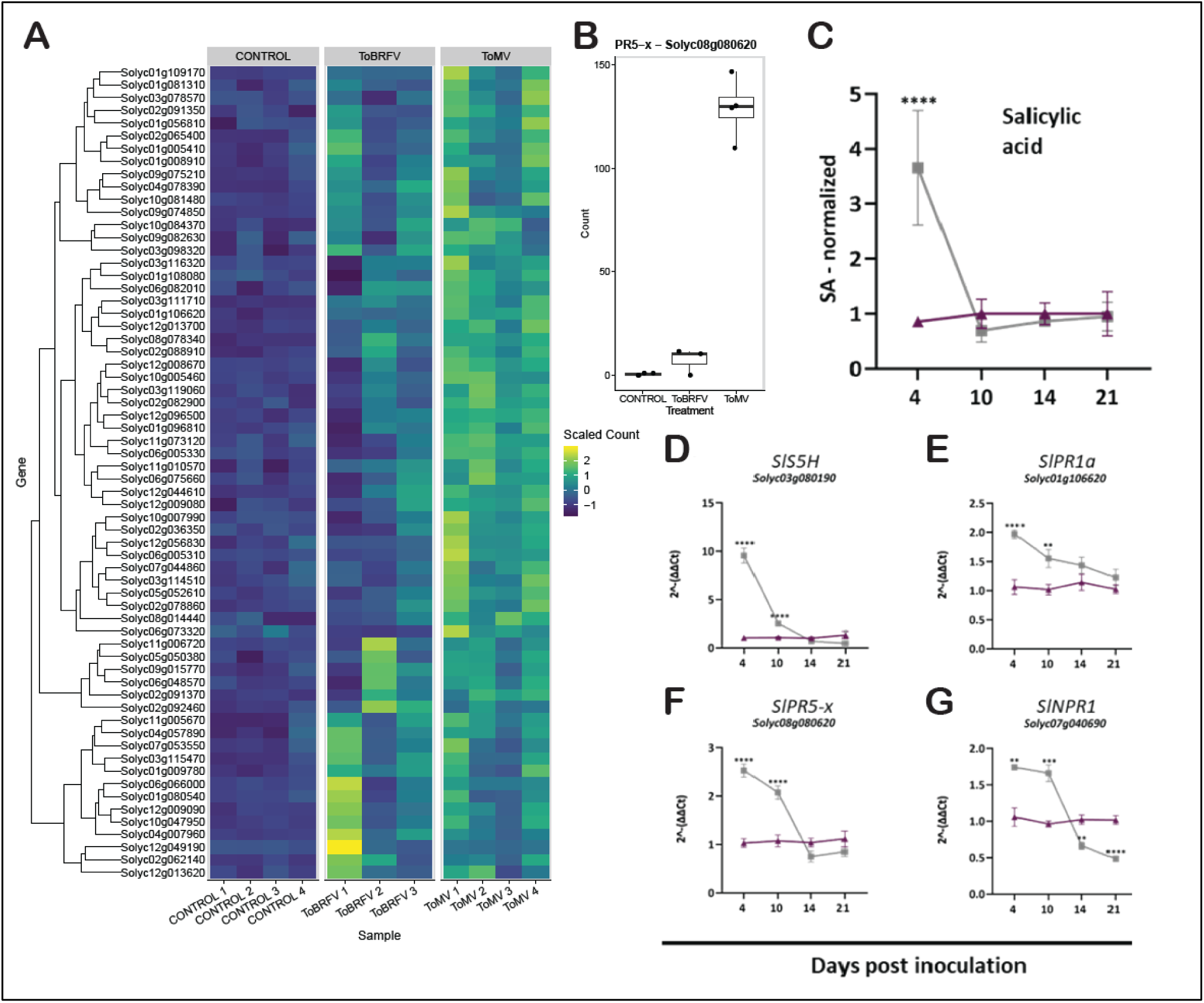
Salicylic acid accumulation and response dynamics in tobamovirus-infected tomato roots. (A) Heat map of salicylic acid (SA) response genes (GO:0009751: response to salicylic acid) upregulated in ToBRFV- or ToMV-infected roots at 10 dpi. Heatmap color indicates the scaled normalized read count. (B) Normalized RNAseq read counts of Pathogenesis Related 5 family gene *PR5-x* (*Solyc08g080620*). (C) Salicylic acid quantification in roots of ToMV-infected tomato plants (cv. Moneymaker) at 4, 10, 14, and 21 days post inoculation (dpi) with ToMV or mock treatment (control). (D–G) Expression of various SA pathway transcripts in ToMV-infected roots: (D) S5H (E) PR1a (F) PR-5x and (G) NPR1. Gene expression was quantified by RT-qPCR using gene-specific primers (Supplementary Table 1). Expression levels were normalized to the geometric mean of the reference genes TIP41 and ACTIN and calculated using the ΔΔCt method. Data represent means ± SE (n = 4–12). Grey and purple points and lines are inoculated and non-inoculated, respectively. Asterisks indicate significant differences between virus-infected samples and the corresponding control (p < 0.05) using one-way ANOVA. *

### Application of Salicylic acid mimics the effect of ToMV over root development

It has been established that SA is a defense hormone, which protects plants against tobamoviruses by reducing their accumulation and restricting their transport (Murphy and Carr, 2002). However, SA is also a root growth regulator that reduces root branching by suppressing lateral root development (Pasternak et al., 2019). To assess the impact of foliar SA application on root development, various concentrations of SA (0.5 mM, 1 mM, and 2 mM) were sprayed on hydroponically grown healthy or ToMV-infected plants. Plants were sprayed with SA every 4-5 days, and their root development and morphology were evaluated at 21 dpi (Fig. 5). Interestingly, application of increasing SA concentrations led to a progressive decline in root biomass (Fig. 5B), total length (Fig. 5C), branch number (Fig. 5D) and branching density (Fig. 5E), in a way that mimics the effect of tobamovirus infection on root development (Fig. 5B-E).

**Figure 5.**
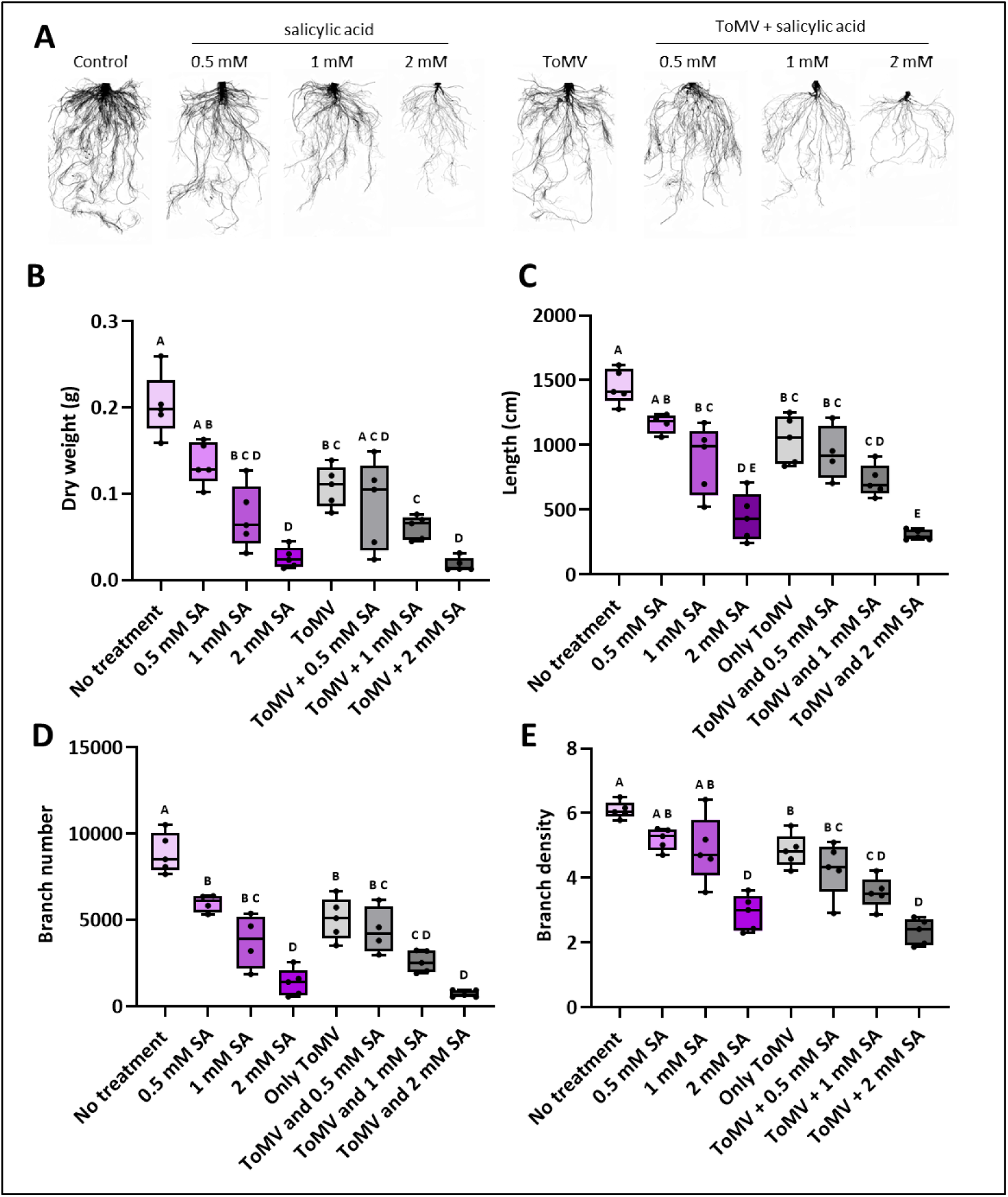
Salicylic acid treatment mimics the effect of ToMV on tomato root development. (A) Representative images of root systems of healthy (green) and ToMV-infected (purple) tomato plants treated with increasing concentrations of SA (0.5, 1, and 2 mM; foliar application) at 21 dpi. (B) Root dry weight (C) total root length (D) branch number (E) branch density. Statistical analysis was performed using one-way ANOVA followed by Tukey’s post hoc test for multiple comparisons or Dunnett’s test when variances were unequal. Bars represent standard error (±SE) (n=5).

In addition, ToMV-infected roots were less sensitive to SA-mediated suppression of root development (Fig. 5D-E), suggesting both ToMV infection and SA act in similar pathways. Application of 2 mM SA resulted in strong reduction in root development that overridden the effect of viral infection on root development (Fig. 5B-D). Since SA treatment can also reduce viral accumulation, viral RNA levels were quantified using RT-qPCR (Supplementary Fig. S3). ToMV RNA was still accumulated in plants treated with 0.5 and 1 mM SA but was reduced in plants treated with 2mM SA (Supplementary Fig. S3). Overall, the results indicate that SA and ToMV infection exert a similar inhibitory effect on root branching. In addition, ToMV infection overrides the effects of lower concentrations of SA. These results suggest that the transient increase in SA accumulation may play a role in tobamovirus-mediated suppression of root branching.

### Tobamovirus infection reprograms the auxin response in tomato roots

Previously, we showed that tobamovirus-mediated root branching suppression is associated with an increase with the expression of the *auxin response factor 10a* (*SlARF10a*), a suppressor of auxin signaling (Vaisman et al., 2022). To gain insights into how auxin signaling is altered during tobamovirus infection, indole-acetic-acid (IAA) levels were quantified in whole roots of tobamovirus-infected plants throughout the course of infection (Supplementary Fig. S4). No changes in IAA levels were observed (Supplementary Fig. S4A), and minimal changes in its breakdown product oxindole-3-acetic acid (ox-IAA; Supplementary Fig. S4B). However, ToMV infected plants (*Solanum lycopersicum* cv. M82) expressing the auxin-responsive reporter *pDR5:VENUS* (Lieberman-Lazarovich et al., 2019) showed a significant increase in auxin signal intensity in the apical root meristem compared with the control (Fig. 6A-B). Thus, it is likely that root auxin sensing and response were affected by infection. Interestingly, most ARFs were down-regulated in tobamovirus-infected roots (Fig. 6C). Within this class, the strongest decrease was observed in auxin response activators, such as *SlARF5,6,7,8*a,8b,*19a* and *19b* (Fig. 6C). Conversely, auxin response suppressor ARFs, such as *SlARF2,3,4* and *10a* were not suppressed, but rather activated (Fig. 6C). Consistent with these findings, the Aux/IAA family of auxin signaling suppressors was generally up-regulated in tobamovirus-infected roots, with the strongest increase in *SlIAA4*, *SlIAA13* and *SlIAA27* (Fig. 6D). To determine if auxin response is altered in tobamovirus-infected roots, the expression of various factors involved in auxin signaling was analyzed (Fig. 6E-J). Consistent with the RNA-seq data, expression of the response activating *SlARF7* and *SlARF19a* was suppressed in ToMV-infected roots, (Fig. 6E-F). In marked contrast, and consistent with our previous findings (Vaisman et al., 2022), auxin-response-suppressive *SlARF4* and *SLARF10a* were up-regulated at 10-14 dpi (Fig. 6G-H). In addition, the auxin signal suppressor *SlIAA4* (Fig. 6I) was upregulated at 10-14 dpi, respectively. In addition, the auxin transporter *SlPIN5* was downregulated at 10 and 21 dpi (Fig. 6J). Collectively, these findings suggest that tobamovirus infection reprograms multiple elements of the auxin response network in tomato roots to potentially suppress root branching.

**Figure 6.**
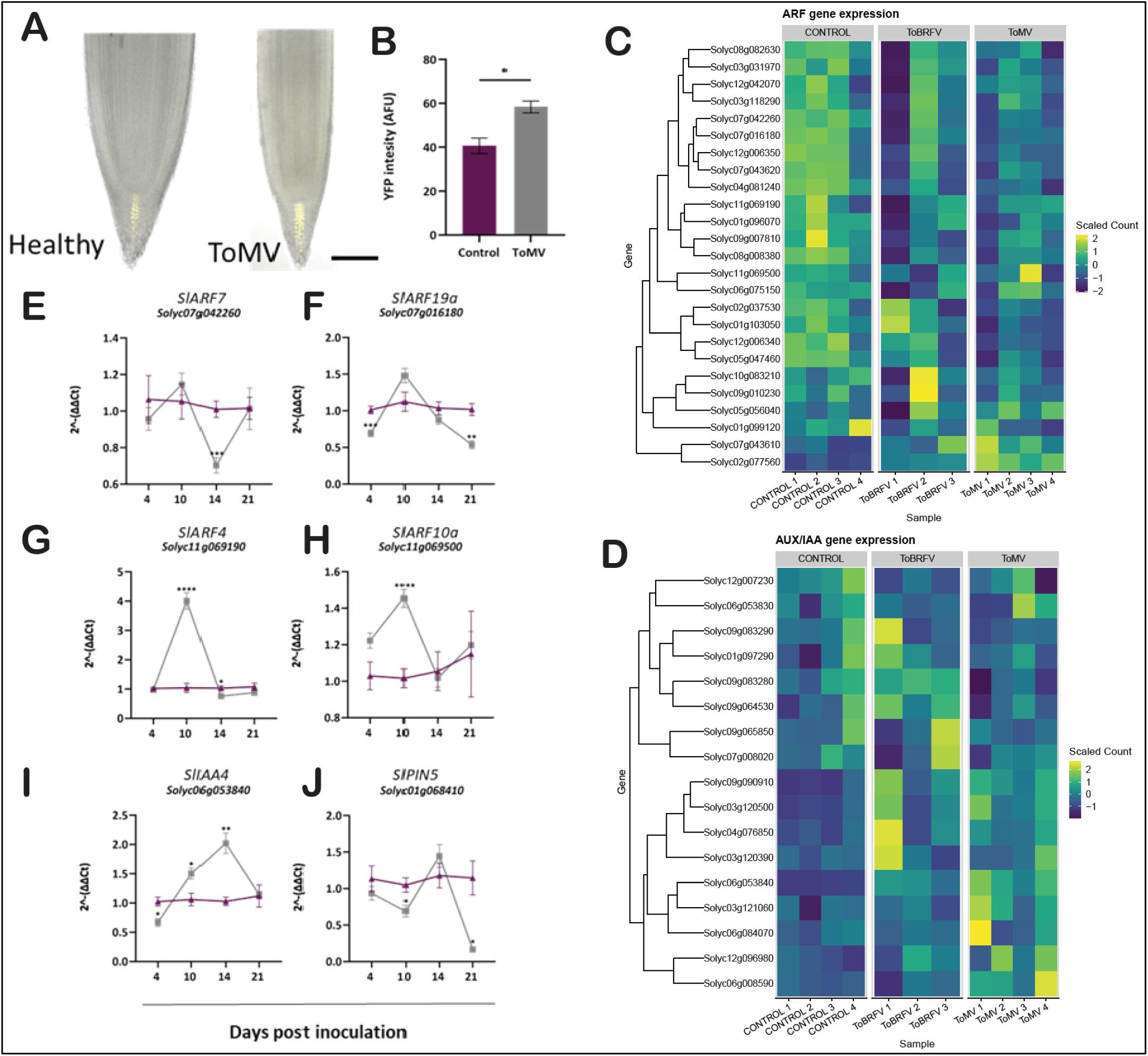
Auxin response dynamics in tobamovirus-infected tomato roots. (A) Representative confocal images of tomato plants (*Solanum lycopersicum* cv. M82) expressing the auxin-responsive reporter p*DR5:VENUS*. Images were taken at 10 days post inoculation (dpi) from. (B) Quantification of VENUS fluorescence intensity in control and ToMV infected roots. (C-D) Heatmaps of auxin response factor (C) and Aux/IAA expression (D) in ToMV and ToBRFV infected roots at 10 dpi. (E-J) Relative expression levels of selected auxin-related genes in roots of tomato plants inoculated with Tomato mosaic virus (ToMV) compared with mock-inoculated controls, at 4-, 10-, 14-, and 21 dpi. Gene expression was quantified using RT-qPCR: (E) *SlARF7* (F) *SlARF19a* (G) *SlARF4* (H) *SlARF10a* (I) *SlIAA4* (J) *SlPIN5*. Expression levels were normalized to the mean of reference genes (*TIP41* and *ACTIN*) and calculated using the ΔΔCt method. Data represent means ± SE of biological replicates (n = 8–12). *p<0.05, **p<0.01 and p<0.001 in unpaired t-test.

### *Auxin Response Factor 19a* regulates tomato root architecture and its response to ToMV infection

In *A. thaliana*, *ARF19* is a key regulator of root development (Okushima et al., 2007; Lee et al., 2009). However, roles of its orthologues in root development of tomato have not yet been determined. Since the expression of *SlARF19a* is reduced during tobamovirus infection (Fig. 6F), we hypothesized that they contribute to the observed changes in root development. To test this, root development was analyzed in *Slarf19a* CRISPR mutants (Israeli et al., 2019; 2023). Plants were grown on hydroponic media and root system architecture was evaluated (Supplementary Fig. S5). Significant reductions were observed in various root development parameters in *Slarf19a* compared to M82 control, including reduction in root length (Supplementary Fig. S5B), root tip number (Supplementary Fig. S5C) and root branch density (Supplementary Fig. S5D).

To further analyze the role of *ARF19a* in root development during tobamovirus infection, an additional experiment was performed in which hydroponically grown mutants were infected with ToMV and root morphology was further analyzed at 14 dpi (Fig. 7A). Mutant plants of another class A ARF, *Slarf8a*, and cv. M82 served as control. In control M82 plants ToMV infection resulted in root biomass reduction of 81.3% (Fig. 7B). However, in *Slarf19a* mutants, root weight was reduced by 26.5% and 43.7% respectively (Fig. 7B), and in *Slarf8a* root weight was reduced by 69.7%. Similarly, in M82 and *slarf8a* plants root length and root tip number were reduced by 83.6% and 81.6% due to ToMV infection (Fig. 7C,D). However, no significant changes in length and tip numbers were observed in ToMV-infected *Slarf19a* mutants, as compared to healthy *Slarf19a* mutants (Fig. 7C). In addition, while ToMV reduced the root branching density of M82 plants by 50.9%, root branching reduction of 33.7%, and 46.2% were observed in ToMV-infected *Slarf19a*, *Slarf8a*, respectively (Fig. 7D). These results suggest that while *SlARF8a* only plays a minor role in root branching and tobamovirus response, *SlARF19a* may play a significant part in these processes.

**Figure 7.**
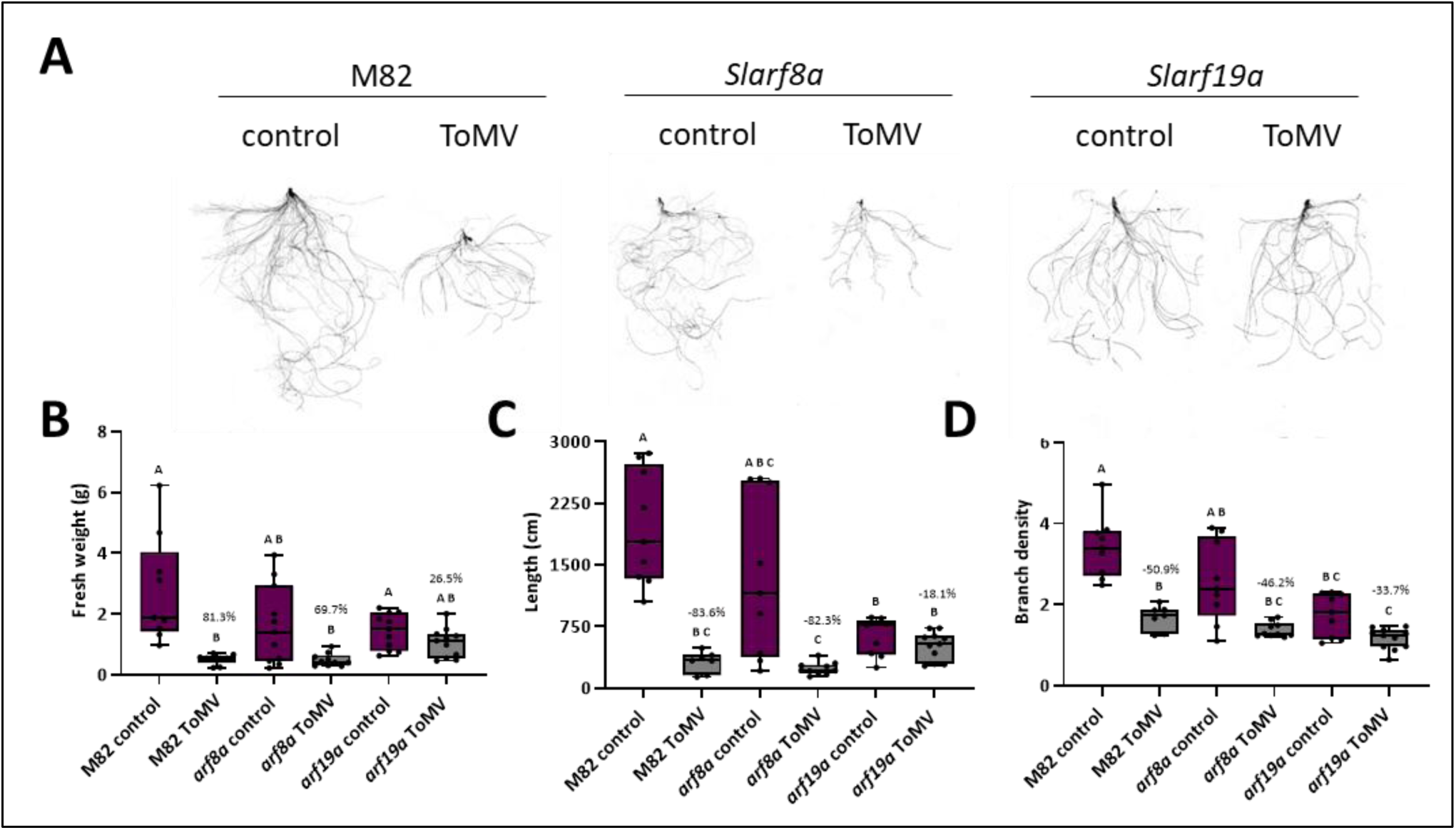
Involvement of *SlARF19a* in tomato root development and response to ToMV. **(A)** Representative scanned root images of healthy and ToMV-infected *Solanum lycopersicum* M82 and *arf19a* and *arf8a mutants* at 14 dpi. **(B–E)** Quantitative analysis of root development parameters: **(B)** fresh root weight, **(C)** total root length, **(D)** Branch density (branch number to total root length). Different letters indicate statistical significance one-way followed multiple-comparison test (p<0.05, n=11).

## Discussion

Tobamoviruses cause substantial damage to various crop plants. While the effects of tobamoviruses on aerial parts of the plants have been expensively explored, little is known about their immune and developmental impacts on root systems. Previously, we identified a substantial developmental response to ToBRFV-infected tomato roots, which predominantly consisted of the suppression of root branching (Vaisman et al., 2022). However, the mechanism by which tobamoviruses affect root development remained to be elucidated. Here, we show that the root developmental response is conserved between the tobamoviruses ToBRFV and ToMV (Figs. 1 and 2). Transcriptional analysis revealed vast reprogramming of the root transcriptome early during infection (Fig. 3). An early induction of SA in the roots of infected plants was subsequently followed by activation of SA-mediated responses during early infection time points (Fig. 4). Exogenous foliar application of SA mimicked the virus’ effect on root development, suggesting a role for SA biosynthesis and activation in tobamovirus mediated suppression of root development. In marked contrast, auxin-mediated development pathways were suppressed in later stages (Fig. 6), including suppression of *SlARF19a* expression, a putative regulator of root development (Fig. 5). Analysis of *Slarf19a* mutants revealed a role for this regulator in during root development and its response to ToMV. Based on our findings, we propose that symptomatic root maldevelopment in response to tobamovirus infection may be mediated in part by reprogramming of auxin pathways by SA-induced defense pathways.

While both ToBRFV and ToMV elicited similar effects over root development, the effect of ToMV was stronger. Consistent with these findings, ToMV infection resulted in a stronger, more consistent impact on the root transcriptome than ToBRFV infection (Fig. 3). The reduced effect of ToBRFV over root development could be due to several factors. First, movement of this virus is attenuated when compared to other tobamoviruses such as TMV and ToMV (Hak and Spiegelman, 2021; Hak et al., 2023). The reduced movement may therefore reduce the rate of viral infection and therefore its transcriptional and developmental responses. While comparison of absolute virus load levels might not directly correlate to symptom severity, in our measurements herein, levels of ToBRFV had not yet peaked in their accumulation by 10 DPI, while those of ToMV had (Fig. 2). Another explanation for the more variable root response to ToBRFV might by that while ToMV is a well-established tomato pathogen which has likely co-evolved with tomato plants for millions of years (Gibbs, 1999), ToBRFV only recently shifted to tomato from another, unknown plant species (Maayan et al., 2018). Thus, ToBRFV may be still in the process of adapting to this new host, and therefore its infection would likely lead to a more variable transcriptional and developmental response. Higher resolution root infection transcriptomes may help indicate whether the reduced ToBRFV response at 10 dpi is due to its slower spread throughout the plant.

Interestingly, almost all DE transcripts in ToBRFV (388/452, 85.8%) were also in DE in ToMV, indicative of a core transcriptional response activated by tobamoviruses in tomato roots. Among the enriched GO categories in both virus-infected roots was ‘response to stimulus’. Homing in on this response in the robust ToMV infection assay, and as evident by GO-term clustering (Fig. 3C) and hormone accumulation (Fig. 4C), the role of SA was clear as a crucial response factor. The function of SA in plant defense responses against viruses has been extensively investigated (Murphy et al., 1999, 2020; Mahmoud et al., 2024). SA inhibits viral replication (Chivasa et al., 1997; Tian et al., 2015), movement (Naylor et al., 1998; Murphy and Carr, 2002; Wang et al., 2013) and invasion into plant meristems (Lee et al., 2016; Incarbone et al., 2023; Hoffman and Incarbone, 2024). In our study, accumulation of SA in the roots was observed very early in disease progression (4 dpi), even before ToMV levels peak (at 10-14 dpi), which subsequently triggers upregulation of multiple SA responsive genes in the roots (Fig. 4). This suggests that SA induction in the roots is likely a rapid and systemic response to viral sensing in the above ground inoculated tissue. Foliar application of SA in tomato has been shown to induce rapid root SA accumulation (Mandal et al., 2009) and perhaps canonical viral induction of SA in the inoculated leaves triggers SA production or transport towards the roots in anticipation of viral movement.

Beyond plant defense, SA is also a known key regulator of shoot and root development (Peng et al., 2021). Activation of SA biosynthesis and signaling often results in plants dwarfism (Zhang et al., 2017; Zeilmaker et al., 2015), while SA downregulation promotes growth and development (Abreu and Munne-Bosch, 2009). Similarly, SA plays a significant role in the suppression of root development (Wang et al., 2007; Pasternak et al., 2019; Tan et al., 2020). Since various viruses induce SA biosynthesis (Yang et al., 2016; Shaw et al., 2019; Murphy et al., 2020), it is logical to hypothesize that SA induction is a symptom determinant for plant viruses. Here, we show that SA-mediated signaling is activated in tobamovirus-infected roots, and that exogenous application of SA mimics the viral effects over root development. In addition, virus-infected plants are less sensitive to SA application, suggesting that the virus and SA acts in the same pathway to suppress root development. These findings suggest that activation of SA may be a determinant of viral infection and potentially the cause of root development symptoms observed in early infection.

The suppressive effect of SA on growth and development can be attributed to its negative regulation of auxin signaling (Wang et al., 2007; Tan et al., 2020; Rawat and Laxmi, 2025). In roots, application of SA interferes with auxin signaling and transport (Pasternak et al., 2019; Kong et al., 2020). Specifically, SA changes auxin distribution in the root apical meristem and alters the expression of several auxin PIN efflux transporters (Pasternak et al., 2019). These changes were associated with suppression of lateral root development. Here, we show that multiple components of the auxin signaling pathway are also altered in tobamovirus-infected roots: while auxin signal activators were generally suppressed, auxin suppressors were activated. While these effects may also be independent from SA, the joint activation of SA-mediated pathways and suppression of auxin signaling pathways may result in the restriction of lateral root development observed in tobamovirus-infected roots. An interesting observation is the upregulation of the auxin-response marker *pDR5:VENUS* in root apical meristems (Fig. 6A-B). It is possible that while auxin response was generally suppressed, local auxin response in root apical meristems was activated. These results are also consistent with the activation of *pDR5* signal observed in *Arabidopsis thaliana* roots treated with moderate concentrations of SA (Pasternak et al. 2019). These results indicate that tobamovirus infection changes the root auxin response in a site-specific manner and potentially in response to early induction of SA.

Manipulation of the SA-auxin signaling network by a plant pathogen was previously observed in *Arabidopsis* plants infected with *Pseudomonas syringae* pv. tomato DC3000 (Kong et al., 2020). In this system, bacterial infection resulted in the up-regulation of *ARF7* and *ARF19*, causing the suppression of SA-mediated lateral root suppression. Interestingly, our findings suggest that tobamovirus act in an opposite manner – SA response is upregulated while *SlARF7* and *SlARF19*a are suppressed. In addition, we find that the function of *SlARF19a* as a regulator of root development is conserved in tomato and Arabidopsis: *Slarf19a* displayed significant reduction in root branching, attenuating response to ToMV infection. These findings establish that *SlARF19a* is critical for root branching in tomato, and that its suppression determines the extent of viral impact on root development.

The significance of tobamovirus-mediated lateral root suppression for viral fitness is still an open question. In a previous study, TMV was shown to be excluded from lateral root meristems (Valentine et al., 2002). It is therefore possible that suppression of lateral roots increases the sites in the root where the virus could be replicated without being suppressed by the plant silencing machinery. In addition, viral infection, including that of tobamoviruses, often confers drought tolerance (Xu et al., 2008; Aguilar et al., 2017; Prakash et al., 2024). It is therefore possible that suppression of lateral root development reduces water uptake by the plant and therefore enables it to survive longer periods of irrigation arrest.

In conclusion, this study finds that tobamovirus-mediated lateral root suppression is a conserved symptom that involves early activation of SA-mediated processes potentially impacting reduction in auxin response. In addition, we find that one of the downregulated auxin response factors, *SlARF19a,* plays a significant role in tomato lateral root development and its response to viral infection. Together, these findings provide insights into the role of plant defense responses and growth-defense tradeoffs in the determination of viral symptoms.

## Data availability

The raw sequence data generated from this project can be retrieved from the NCBI SRA database under BioSample SAMNXXXX/PRJNXXXXXXXX. Code for producing figures for the manuscript is available at github.com/bmansfeld/tomato_virus_roots/.

## Supplementary data

**Supplementary Fig. S1.** Accumulation of ToMV and ToBRFV in tomato plants.

**Supplementary Fig. S2.** Hydroponic growing system allows detailed analysis of root growth under viral infection.

**Supplementary Fig. S3.** ToMV RNA accumulation in plants treated with salicylic acid.

**Supplementary Fig. S4.** Quantification of IAA and oxidized IAA in tobamovirus-infected roots.

**Supplementary Fig. S5.** *Auxin response factor 19a* (*ARF19a*) regulates tomato root development.

**Supplementary table S1.** Oligonucleotides used in this manuscript.

**Supplementary table S2.** Differential expression analysis in ToMV- and ToBRFV-infected roots.

**Supplementary table S3.** Gene ontology enrichment analysis of transcripts upregulated in ToMV- and ToBRFV-infected roots.

**Supplementary table S4.** Gene ontology enrichment analysis of transcripts downregulated in ToMV- and ToBRFV-infected roots.

## Acknowledgements

We thank Prof. Naomi Ori from the Hebrew University of Jerusalem for kindly sharing tomato *Slarf8a* and *Slarf19a* mutant seeds. This work was supported by the Israel Science Foundation (GA No. 3347/25) and German Israeli Foundation for Scientific Research and Development (I-1543-500.12/2021).

## Author contributions

D.W., B.N.M and Z.S. designed the research; D.W., F.S and M.W-C conducted the experiments; D.W. and B.N.M conducted the RNA data analysis. D.W, B.N.M and Z.S. and wrote the manuscript.

## References

Abreu, M. E., & Munne-Bosch, S. (2009). Salicylic acid deficiency in NahG transgenic lines and sid2 mutants increases seed yield in the annual plant Arabidopsis thaliana. Journal of experimental botany, 60(4), 1261–1271.

Adams, M. J., Adkins, S., Bragard, C., Gilmer, D., Li, D., MacFarlane, S. A., Wong, S. M., Melcher, U., Ratti, C., Ryu, K. H., & ICTV Report Consortium. (2017). ICTV virus taxonomy profile: Virgaviridae. Journal of General Virology, 98(8), 1999–2000.

Aguilar, E., Cutrona, C., Del Toro, F. J., Vallarino, J. G., Osorio, S., Pérez-Bueno, M. L., … & Tenllado, F. (2017). Virulence determines beneficial trade-offs in the response of virus-infected plants to drought via induction of salicylic acid. Plant, Cell & Environment, 40(12), 2909–2930.

Alexa, A., Rahnenführer, J., & Lengauer, T. (2006). Improved scoring of functional groups from gene expression data by decorrelating GO graph structure. Bioinformatics, 22(13), 1600–1607.

Bagautdinova, Z. Z., Omelyanchuk, N., Tyapkin, A. V., Kovrizhnykh, V. V., Lavrekha, V. V., & Zemlyanskaya, E. V. (2022). Salicylic acid in root growth and development. International Journal of Molecular Sciences, 23(4), 2228.

Baulcombe, D. C. (2015). VIGS, HIGS and FIGS: small RNA silencing in the interactions of viruses or filamentous organisms with their plant hosts. Current opinion in plant biology, 26, 141–146.

Chellappan, P., Vanitharani, R., & Fauquet, C. M. (2005). MicroRNA-binding viral protein interferes with Arabidopsis development. Proceedings of the National Academy of Sciences, 102(29), 10381–10386.

Chivasa, S., Murphy, A. M., Naylor, M., & Carr, J. P. (1997). Salicylic acid interferes with tobacco mosaic virus replication via a novel salicylhydroxamic acid-sensitive mechanism. The Plant Cell, 9(4), 547–557.

Csorba, T., Kontra, L., & Burgyán, J. (2015). Viral silencing suppressors: Tools forged to fine-tune host–pathogen coexistence. Virology, 479–480, 85–103.

Gao, M., & Lozano-Durán, R. (2025). Symptom development in plant viral diseases: What, how, and why? Annual Review of Phytopathology, 63.

Gao, M., Aguilar, E., Gómez, B. G., Medina-Puche, L., Fan, P., Ontiveros, I., Pan, S., Tan, H., Wei, H., von Roepenack-Lahaye, E., & Chen, N. (2025). A plant virus causes symptoms through the deployment of a host-mimicking protein domain to attract the insect vector. Molecular Plant, 18(6), 1029–1046.

Gerola, F. M., Bassi, M., Favali, M. A., & Betto, E. (1969). An electron microscopy study of the penetration of tobacco mosaic virus into leaves following experimental inoculation. Virology, 38(3), 380–386.

Gibbs, A. (1999). Evolution and origins of tobamoviruses. Philosophical Transactions of the Royal Society of London. Series B: Biological Sciences, 354(1383), 593–602.

Gnanasekaran, P., Zhai, Y., Kamal, H., Smertenko, A., & Pappu, H. R. (2023). A plant virus protein, NIa-pro, interacts with indole-3-acetic acid-amido synthetase, whose levels positively correlate with disease severity. Frontiers in Plant Science, 14, 1112821.

Gosalves-Bernal, B., Genoves, A., Navarro, J. A., Pallas, V., & Sanchez-Pina, M. A. (2008). Distribution and pathway for phloem-dependent movement of Melon necrotic spot virus in melon plants. Molecular Plant Pathology, 9(4), 447–461.

Hak, H., & Spiegelman, Z. (2021). The tomato brown rugose fruit virus movement protein overcomes Tm-22 resistance in tomato while attenuating viral transport. Molecular Plant-Microbe Interactions, 34(9), 1024–1032.

Hak, H., Raanan, H., Schwarz, S., Sherman, Y., Dinesh-Kumar, S. P., & Spiegelman, Z. (2023). Activation of Tm-22 resistance is mediated by a conserved cysteine essential for tobacco mosaic virus movement. Molecular Plant Pathology, 24(8), 838–848.

Heinlein, M. (2015). Plant virus replication and movement. Virology, 479, 657–671.

Hendelman, A., Kravchik, M., Stav, R., Zik, M., Lugassi, N., & Arazi, T. (2013). The developmental outcomes of P0-mediated ARGONAUTE destabilization in tomato. Planta, 237(1), 363–377.

Hoffmann, G., & Incarbone, M. (2024). A resilient bunch: stem cell antiviral immunity in plants. New Phytologist, 241(4), 1415–1420.

Hosmani, P. S., Flores-Gonzalez, M., van de Geest, H., Maumus, F., Bakker, L. V., Schijlen, E., van Haarst, J., Cordewener, J., Sanchez-Perez, G., Peters, S., & Fei, Z. (2019). An improved de novo assembly and annotation of the tomato reference genome using single-molecule sequencing, Hi-C proximity ligation and optical maps. bioRxiv, 767764.

Ibrahim, A., Sasaki, N., Schoelz, J. E., & Nelson, R. S. (2025). Tobacco mosaic virus movement: From capsid disassembly to transport through plasmodesmata. Viruses, 17(2), 214.

Inaba, J.-I., Kim, B. M., Shimura, H., & Masuta, C. (2011). Virus-induced necrosis is a consequence of direct interaction between a viral RNA-silencing suppressor and a host catalase. Plant Physiology, 156(4), 2026–2036.

Incarbone, M., Bradamante, G., Pruckner, F., Wegscheider, T., Rozhon, W., Nguyen, V., … & Scheid, O. M. (2023). Salicylic acid and RNA interference mediate antiviral immunity of plant stem cells. Proceedings of the National Academy of Sciences, 120(42), e2302069120.

Israeli, A., Capua, Y., Shwartz, I., Tal, L., Meir, Z., Levy, M., Bar, M., Efroni, I., & Ori, N. (2019). Multiple auxin-response regulators enable stability and variability in leaf development. Current Biology, 29(11), 1746–1759.

Israeli, A., Schubert, R., Man, N., Teboul, N., Serrani Yarce, J. C., Rosowski, E. E., Wu, M.-F., Levy, M., Efroni, I., Ljung, K., & Hause, B. (2023). Modulating auxin response stabilizes tomato fruit set. Plant Physiology, 192(3), 2336–2355.

Ishibashi, K., & Ishikawa, M. (2016). Replication of tobamovirus RNA. Annual Review of Phytopathology, 54(1), 55–78.

Ishibashi, K., Kubota, K., Kano, A., & Ishikawa, M. (2023). Tobamoviruses: old and new threats to tomato cultivation. Journal of General Plant Pathology, 89(6), 305–321.

Jones, P., Binns, D., Chang, H.-Y., Fraser, M., Li, W., McAnulla, C., McWilliam, H., Maslen, J., Mitchell, A., Nuka, G., & Pesseat, S. (2014). InterProScan 5: genome-scale protein function classification. Bioinformatics, 30(9), 1236–1240.

Jones, R. A. (2021). Global plant virus disease pandemics and epidemics. Plants, 10(2), 233.

Jones, R. A., & Naidu, R. A. (2019). Global dimensions of plant virus diseases: current status and future perspectives. Annual Review of Virology, 6(1), 387–409.

Kan, Y., & Citovsky, V. (2025). The roles of movement and coat proteins in the transport of tobamoviruses between plant cells. Frontiers in Plant Science, 16, 1580554.

Karlova, R., Boer, D., Hayes, S., & Testerink, C. (2021). Root plasticity under abiotic stress. Plant Physiology, 187(3), 1057–1070.

Kasschau, K. D., Xie, Z., Allen, E., Llave, C., Chapman, E. J., Krizan, K. A., & Carrington, J. C. (2003). P1/HC-Pro, a viral suppressor of RNA silencing, interferes with Arabidopsis development and miRNA function. Developmental Cell, 4(2), 205–217.

Kehr, J., Morris, R.J. and Kragler, F., 2022. Long-distance transported RNAs: from identity to function. Annual review of plant biology, 73(1), 457–474.

Kong, X., Zhang, C., Zheng, H., Sun, M., Zhang, F., Zhang, M., … & Ding, Z. (2020). Antagonistic interaction between auxin and SA signaling pathways regulates bacterial infection through lateral root in Arabidopsis. Cell Reports, 32(8).

Kubota, K., Tsuda, S., Tamai, A., & Meshi, T. (2003). Tomato mosaic virus replication protein suppresses virus-targeted post-transcriptional gene silencing. Journal of Virology, 77(20), 11016–11026.

Lacerda, A. L., Fonseca, L. N., Blawid, R., Boiteux, L. S., Ribeiro, S. G., & Brasileiro, A. C. (2015). Reference gene selection for qPCR analysis in tomato–begomovirus interaction. PLOS ONE, 10(8), e0136820.

Lee, H. W., Kim, N. Y., Lee, D. J., & Kim, J. (2009). LBD18/ASL20 regulates lateral root formation in combination with LBD16/ASL18 downstream of ARF7 and ARF19 in Arabidopsis. Plant physiology, 151(3), 1377–1389.

Lee, W. S., Fu, S. F., Li, Z., Murphy, A. M., Dobson, E. A., Garland, L., … & Carr, J. P. (2016). Salicylic acid treatment and expression of an RNA-dependent RNA polymerase 1 transgene inhibit lethal symptoms and meristem invasion during tobacco mosaic virus infection in Nicotiana benthamiana. BMC plant biology, 16(1), 15.

Liebman-Lazarovich, M., Yahav, C., Israeli, A., & Efroni, I. (2019). Deep conservation of cis-element variants regulating plant hormonal responses. The Plant Cell, 31(11), 2559–2572.

Li, A., Sun, X., & Liu, L. (2022). Action of salicylic acid on plant growth. Frontiers in Plant Science, 13, 878076.

Livak, K. J., & Schmittgen, T. D. (2001). Analysis of relative gene expression data using real-time quantitative PCR and the 2⁻ΔΔCT method. Methods, 25(4), 402–408.

Love, M. I., Huber, W., & Anders, S. (2014). Moderated estimation of fold change and dispersion for RNA-seq data with DESeq2. Genome Biology, 15(12), 550.

Lunello, P., Mansilla, C., Sánchez, F., & Ponz, F. (2007). A developmentally linked, transient loss of virus from roots of Arabidopsis thaliana. Molecular Plant-Microbe Interactions, 20(12), 1589–1595.

Luria, N., Smith, E., Reingold, V., Bekelman, I., Lapidot, M., Levin, I., Elad, N., Tam, Y., Sela, N., Abu-Ras, A., & Ezra, N. (2017). A new Israeli tobamovirus isolate infects tomato plants harboring Tm-22. PLOS ONE, 12(1), e0170429.

Maayan, Y., Pandaranayaka, E. P., Srivastava, D. A., Lapidot, M., Levin, I., Dombrovsky, A., & Harel, A. (2018). Using genomic analysis to identify tomato Tm-2 resistance-breaking mutations and their underlying evolutionary path in a new and emerging tobamovirus. Archives of virology, 163(7), 1863–1875.

Malamy, J., et al. (1990). Salicylic acid: a likely endogenous signal in the resistance response of tobacco to viral infection. Science, 250(4983), 1002–1004.

Mahmood, M. A., Naqvi, R. Z., Amin, I., & Mansoor, S. (2024). Salicylic acid-driven innate antiviral immunity in plants. Trends in Plant Science, 29(7), 715–717.

Mandal, S., Mallick, N., & Mitra, A. (2009). Salicylic acid-induced resistance to Fusarium oxysporum f. sp. lycopersici in tomato. Plant physiology and Biochemistry, 47(7), 642–649.

Marash, I., Leibman-Markus, M., Gupta, R., Israeli, A., Teboul, N., Avni, A., Ori, N., & Bar, M. (2024). Abolishing ARF8A activity promotes disease resistance in tomato. Plant Science, 343, 112064.

Muellender, M. M., Savenkov, E. I., Reichelt, M., Varrelmann, M., & Liebe, S. (2022). The virulence factor p25 interacts with multiple Aux/IAA proteins. Frontiers in Microbiology, 12, 809690.

Murphy, A. M., Chivasa, S., Singh, D. P., & Carr, J. P. (1999). Salicylic acid-induced resistance to viruses and other pathogens: a parting of the ways?. Trends in plant science, 4(4), 155–160.

Murphy, A. M., Zhou, T., & Carr, J. P. (2020). An update on salicylic acid biosynthesis, its induction, and potential exploitation by plant viruses. Current Opinion in Virology, 42, 8–17.

Murphy, A. M., & Carr, J. P. (2002). Salicylic acid has cell-specific effects on TMV replication. Plant Physiology, 128(2), 552–563.

Naylor, M., Murphy, A. M., Berry, J. O., & Carr, J. P. (1998). Salicylic acid can induce resistance to plant virus movement. Molecular Plant-Microbe Interactions, 11(9), 860–868.

Okushima, Y., Fukaki, H., Onoda, M., Theologis, A., & Tasaka, M. (2007). ARF7 and ARF19 regulate lateral root formation via direct activation of LBD/ASL genes. The Plant Cell, 19(1), 118–130.

Pallas, V., & García, J. A. (2011). How do plant viruses induce disease? Journal of General Virology, 92(12), 2691–2705.

Pang, W., Crow, W. T., Luc, J. E., McSorley, R., Giblin-Davis, R. M., Kenworthy, K. E., & Kruse, J. K. (2011). Comparison of water displacement and WinRHIZO software for root analysis. Plant Disease, 95(10), 1308–1310.

Panno, S., Davino, S., Caruso, A. G., Bertacca, S., Crnogorac, A., Mandić, A., Noris, E., & Matić, S. (2021). Common tomato diseases in the Mediterranean basin. Agronomy, 11(11), 2188.

Pasternak, T., Groot, E. P., Kazantsev, F. V., Teale, W., Omelyanchuk, N., Kovrizhnykh, V., … & Mironova, V. V. (2019). Salicylic acid affects root meristem patterning via auxin distribution in a concentration-dependent manner. Plant physiology, 180(3), 1725–1739.

Patro, R., Duggal, G., Love, M. I., Irizarry, R. A., & Kingsford, C. (2017). Salmon: fast and bias-aware transcript quantification. Nature Methods, 14(4), 417–419.

Prakash, V., Sharma, V., Devendran, R., Prajapati, R., Ahmad, B., & Kumar, R. (2024). A transition from enemies to allies: how viruses improve drought resilience in plants. Stress Biology, 4(1), 33.

Peltier, C., Schmidlin, L., Klein, E., Taconnat, L., Prinsen, E., Erhardt, M., Heintz, D., Weyens, G., Lefebvre, M., Renou, J.-P., & Gilmer, D. (2011). BNYYV p25 induces hormonal changes and root branching. Transgenic Research, 20, 443–466.

Peng, Y., Yang, J., Li, X., & Zhang, Y. (2021). Salicylic acid: biosynthesis and signaling. Annual review of plant biology, 72(1), 761–791.

Pokotylo, I., Hodges, M., Kravets, V., & Ruelland, E. (2022). A ménage à trois: salicylic acid, growth inhibition, and immunity. Trends in Plant Science, 27(5), 460–471.

Rawat, S. S., & Laxmi, A. (2025). Rooted in communication: exploring auxin–salicylic acid nexus in root growth and development. Plant, Cell & Environment, 48(6), 4140–4160.

Salem, N. M., Jewehan, A., Aranda, M. A., & Fox, A. (2023). Tomato brown rugose fruit virus pandemic. Annual Review of Phytopathology, 61(1), 137–164.

Samuel, G. (1934). The movement of tobacco mosaic virus within the plant. Annals of Applied Biology, 21(1), 90–111.

Sayols, S. (2023). rrvgo: a Bioconductor package for interpreting lists of Gene Ontology terms. microPublication biology, 2023, 10–17912.

Schneider, H. M., & Lynch, J. P. (2020). Should root plasticity be a crop breeding target? Frontiers in Plant Science, 11, 546.

Shaw, J., Yu, C., Makhotenko, A. V., Makarova, S. S., Love, A. J., Kalinina, N. O., … & Taliansky, M. E. (2019). Interaction of a plant virus protein with the signature Cajal body protein coilin facilitates salicylic acid-mediated plant defence responses. New Phytologist, 224(1), 439–453.

Shihan, M. H., Novo, S. G., Le Marchand, S. J., Wang, Y., & Duncan, M. K. (2021). A simple method for quantitating confocal fluorescent images. Biochemistry and biophysics reports, 25, 100916.

Singh, D. P., Moore, C. A., Gilliland, A., & Carr, J. P. (2004). Activation of antiviral defence mechanisms by salicylic acid. Molecular Plant Pathology, 5(1), 57–63.

Soneson, C., Love, M. I., & Robinson, M. D. (2016). Differential analyses for RNA-seq. F1000Research, 4, 1521.

Spiegelman, Z., & Dinesh-Kumar, S. P. (2023). Breaking boundaries: the perpetual interplay between tobamoviruses and plant immunity. Annual Review of Virology, 10(1), 455–476.

Tan, S., Abas, M., Verstraeten, I., Glanc, M., Molnár, G., Hajný, J., Lasák, P., Petřík, I., Russinova, E., Petrášek, J., & Novák, O. (2020). Salicylic acid targets PP2A to attenuate growth in plants. Current Biology, 30(3), 381–395.

Tatineni, S., & Hein, G. L. (2023). Plant viruses of agricultural importance. Phytopathology, 113(2), 117–141.

Thiel, H., & Varrelmann, M. (2009). BNYYV P25 pathogenicity factor–interacting proteins in sugar beet. Molecular Plant-Microbe Interactions, 22(8), 999–1010.

Tian, M., Sasvari, Z., Gonzalez, P. A., Friso, G., Rowland, E., Liu, X. M., … & Klessig, D. F. (2015). Salicylic acid inhibits the replication of tomato bushy stunt virus by directly targeting a host component in the replication complex. Molecular Plant-Microbe Interactions, 28(4), 379–386.

Trapnell, C., Roberts, A., Goff, L., Pertea, G., Kim, D., Kelley, D. R., Pimentel, H., Salzberg, S. L., Rinn, J. L., & Pachter, L. (2012). Differential gene and transcript expression analysis with TopHat and Cufflinks. Nature Protocols, 7(3), 562–578.

Turgeon, R., & Wolf, S. (2009). Phloem transport: cellular pathways and molecular trafficking. Annual review of plant biology, 60, 207–221.

Vaisman, M., Hak, H., Arazi, T., & Spiegelman, Z. (2022). The impact of tobamovirus infection on root development involves induction of auxin response factor 10a in tomato. Plant and Cell Physiology, 63(12), 1980–1993.

Valentine, T. A., Roberts, I. M., & Oparka, K. J. (2002). Inhibition of TMV replication in lateral roots depends on an activated meristem-derived signal. Protoplasma, 219(3–4), 184–196.

Villordon, A. Q., & Clark, C. A. (2014). Variation in virus symptoms and root architecture during storage root initiation. PLOS ONE, 9(9), e107384.

Wang, X., Sager, R., Cui, W., Zhang, C., Lu, H., & Lee, J. Y. (2013). Salicylic acid regulates plasmodesmata closure during innate immune responses in Arabidopsis. The Plant Cell, 25(6), 2315–2329.

Wickham, H., Averick, M., Bryan, J., Chang, W., McGowan, L. D., François, R., Grolemund, G., Hayes, A., Henry, L., Hester, J., & Kuhn, M. (2019). Welcome to the tidyverse. Journal of Open Source Software, 4(43), 1686.

Wimalanathan, K., & Lawrence-Dill, C. J. (2021). Gene ontology meta annotator for plants (GOMAP). Plant Methods, 17(1), 54.

Xu, P., Chen, F., Mannas, J. P., Feldman, T., Sumner, L. W., & Roossinck, M. J. (2008). Virus infection improves drought tolerance. New Phytologist, 180(4), 911–921.

Yadav, S., & Chhibbar, A. K. (2018). Plant–virus interactions. In Molecular Aspects of Plant–Pathogen Interaction (pp. 43–77). Springer.

Yang, L., Xu, Y., Liu, Y., Meng, D., Jin, T., & Zhou, X. (2016). HC-Pro viral suppressor from tobacco vein banding mosaic virus interferes with DNA methylation and activates the salicylic acid pathway. Virology, 497, 244–250.

Zamfir, A. D., Babalola, B. M., Fraile, A., McLeish, M. J., & García-Arenal, F. (2023). Tobamoviruses show broad host ranges and little genetic diversity across habitats. Phytopathology, 113(9), 1697–1707.

Zhang, S., Griffiths, J. S., Marchand, G., Bernards, M. A., & Wang, A. (2022). Tomato brown rugose fruit virus: an emerging RNA virus threatening tomato production worldwide. Molecular Plant Pathology, 23(9), 1262–1277.

Zhang, Y., Zhao, L., Zhao, J., Li, Y., Wang, J., Guo, R., Gan, S., Liu, C.J. and Zhang, K., 2017. S5H/DMR6 encodes a salicylic acid 5-hydroxylase that fine-tunes salicylic acid homeostasis. Plant Physiology, 175(3), 1082–1093.

Zeilmaker, T., Ludwig, N. R., Elberse, J., Seidl, M. F., Berke, L., Van Doorn, A., … & Van den Ackerveken, G. (2015). DOWNY MILDEW RESISTANT 6 and DMR 6-LIKE OXYGENASE 1 are partially redundant but distinct suppressors of immunity in Arabidopsis. The Plant Journal, 81(2), 210–222.

Zhu, A., Ibrahim, J. G., & Love, M. I. (2019). Heavy-tailed priors for sequence count data. Bioinformatics, 35(12), 2084–2092.

