## Supplementary Fig. for "Antagonism between salicylic acid and auxin responses directs root development during *Tobamovirus* infection"

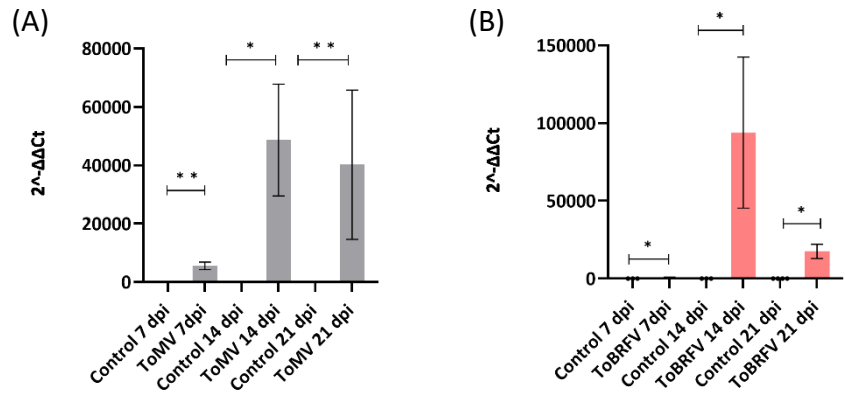

**Supplementary Fig. S1. Accumulation of ToMV and ToBRFV in tomato plants.** RT-qPCR quantification of ToMV (A) and ToBRFV (B) accumulation from RNA extracted from young leaves of infected plants, normalized to the housekeeping transcript *TIP41*. statistical significance was assessed using Student's *t*-test with Welch's correction. Data are presented as mean ± SEM. \*P-value < 0.05, \*\*P-value < 0.01, n= 6-7.

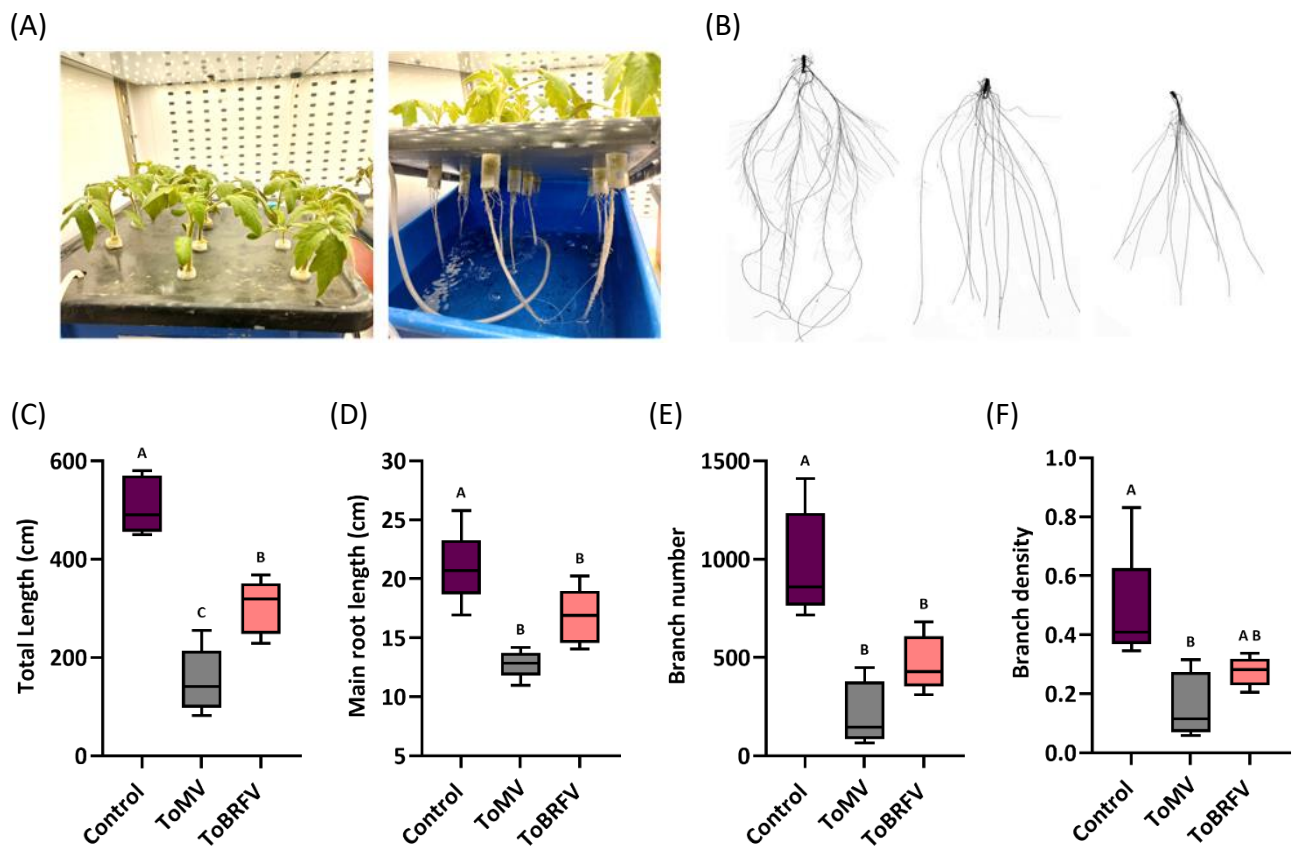

**Supplementary Fig. S2. Hydroponic growing system allows detailed analysis of root growth under viral infection.** (A) Tomato plants grown in hydroponics allow non-destructive analysis of roots. (B) Winrhizo root scans of healthy, ToBRFV- and ToMV-infected plants, showing marked suppression of root branching. (C-F) Quantification of total root length (C), primary root length (D), number of root branches (E) and root branching density (F). Different letters indicate significance in Tukey-HSD test.

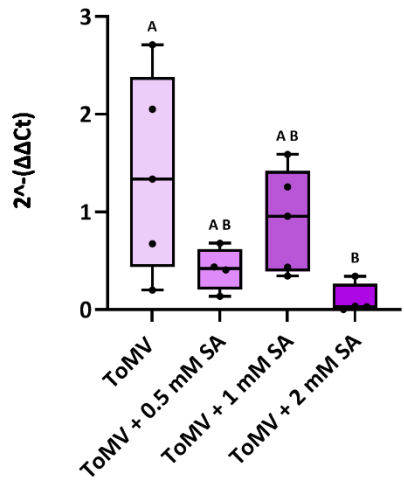

**Supplementary Fig. S3. ToMV RNA accumulation in plants treated with salicylic acid.**

Levels of ToMV RNA were quantified in young leaves of *Solanum lycopersicum* (cv. Moneymaker) plants infected with ToMV and treated with increasing concentrations of salicylic acid (SA; 0.5, 1, and 2 mM; foliar application). Samples were collected at 21 dpi. Statistical analysis was performed using the Kruskal–Wallis non-parametric test followed by Dunn’s multiple-comparison test. Different letter indicate statistical significance ($P < 0.05$) ($n = 5$).

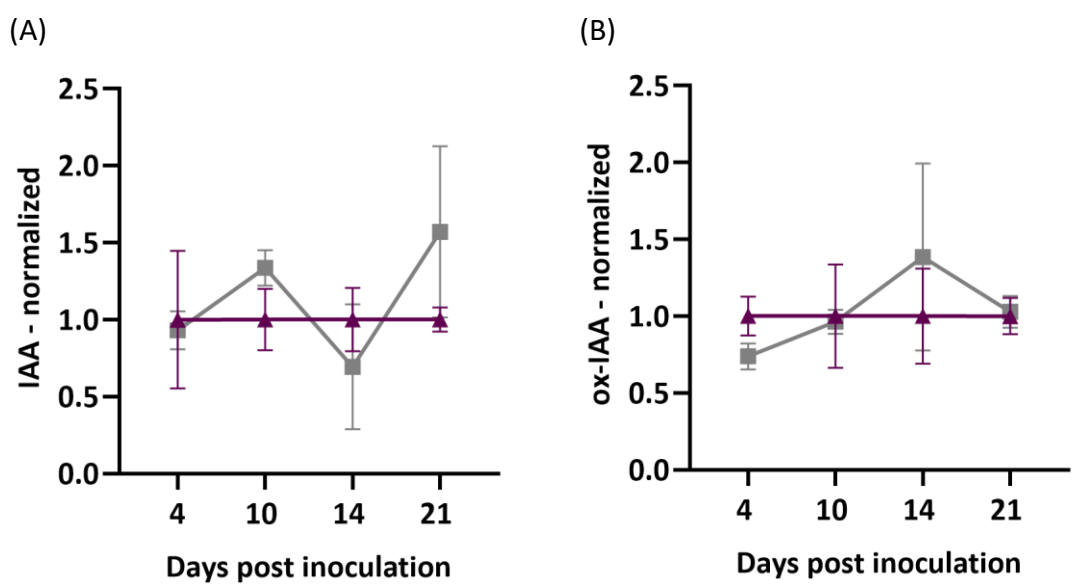

**Supplementary Fig. S4. Quantification of IAA and oxidized IAA in tobamovirus-infected roots.** Hormone quantification analyses of tomato seedling roots (cv. Moneymaker) at 4, 14, and 21 days post inoculation (dpi) with ToMV or mock treatment (control) **(A)** IAA and **(B)** ox-IAA quantification by LC–MS. Hormone levels were normalized to the fresh weight of the extracted tissue ($\text{mg g}^{-1}$ FW) and further normalized to the mean of the corresponding control treatment at each time point. Four biological replicates were analyzed per treatment, each consisting of roots pooled from 2–3 plants. Bars represent standard deviation (SD). and statistical significance was determined using one-way ANOVA followed by Tukey’s multiple-comparison test.

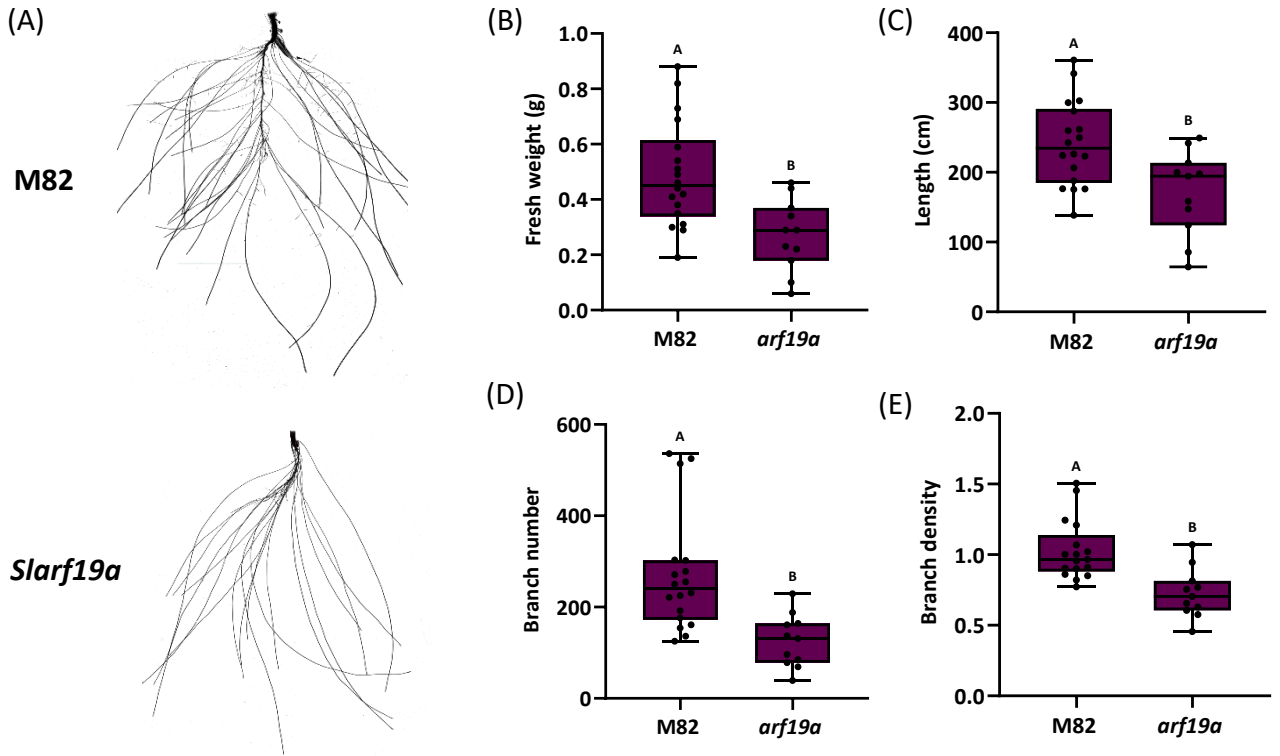

**Supplementary Fig. S5. *Auxin response factor 19a* (*SlARF19a*) regulates tomato root development.** (A) Representative image of M82 and Slarf19a root system (B–D) Quantitative analysis of root development parameters in *Solanum lycopersicum* (cv. M82) and *Slarf19a* mutants grown for 14 days on hydroponic system: (B) fresh root weight (C) total root length (D) branch number (E) branch density (ratio of branch number to total length). Statistical significance was determined using one-way ANOVA followed by Tukey's or Dunnett's multiple-comparison test. Different letters indicate statistical significance (n = 11-18).
